# Msp1 and Pex19-Pex3 cooperate to achieve correct localization of Pex15 to peroxisomes

**DOI:** 10.1101/2024.12.11.627694

**Authors:** Shunsuke Matsumoto, Yoshiki Kogure, Suzuka Ono, Tomoyuki Numata, Toshiya Endo

**Affiliations:** Department of Bioscience and Biotechnology, Graduate School of Bioresource and Bioenvironmental Sciences, Kyushu University, 744 Motooka, Nishi-ku, Fukuoka, 819-0395, Japan; Faculty of Life Sciences, Kyoto Sangyo University, Kamigamo-Motoyama, Kita-ku, Kyoto 603-8555, Japan; Institute for Protein Dynamics, Kyoto Sangyo University, Kamigamo-Motoyama, Kita-ku, Kyoto 603-8555, Japan

## Abstract

Yeast Msp1 is a dual-localized AAA-ATPase on the mitochondrial outer membrane (OM) and peroxisomal membrane. We previously showed that Msp1 transfers mistargeted tail-anchored (TA) proteins from mitochondria to the endoplasmic reticulum (ER) for degradation or delivery to their original destinations. However, the mechanism by which Msp1 in mitochondria and peroxisomes handles authentic peroxisomal TA proteins remains unclear. We show that newly synthesized Pex15 is targeted to peroxisomes primarily *via* the Pex19- and Pex3-dependent pathway. Mistargeted Pex15 on the mitochondrial OM is extracted by mitochondrial Msp1, and transferred to the ER *via* the guided-entry of TA proteins pathway for degradation or to peroxisomes *via* the Pex19-Pex3 pathway. Intriguingly, endogenous Pex15 localized in peroxisomes is also extracted from the membranes by peroxisomal Msp1, but returns to peroxisomes *via* the Pex19-Pex3 pathway. These results suggest that correct Pex15 localization to peroxisomes relies on not only the initial targeting by Pex19-Pex3 but also the constant re-routing by Msp1 and Pex19-Pex3.

## Introduction

The correct delivery of proteins to their intended organelle is essential for maintaining normal cellular functions. In eukaryotic cells, approximately one-fourth of the proteome is composed of membrane proteins (Hegde and Keenan, 2022). Tail-anchored (TA) proteins are a distinct class of membrane proteins characterized by a single transmembrane (TM) segment located near the C-terminus (Wattenberg and Lithgow, 2001). They are estimated to comprise 3%–5% of the total membrane proteome and play vital roles in a variety of cellular processes, including membrane fusion, protein translocation, apoptosis, and autophagy (Chio et al., 2017; Wang and Walter, 2020). TA proteins are anchored to organelle membranes by their C-terminal TM segment, with the N-terminal region facing the cytosolic side. The targeting information for TA proteins is defined by the hydrophobicity and length of the TM segment, along with the positively charged amino acid residues near the TM segment (Borgese and Fasana, 2011).

TA proteins are targeted to organelle membranes in a post-translational manner, since their targeting sequence remains hidden within the ribosomal tunnel until translation is complete (Borgese and Fasana, 2011). After synthesis, TA proteins are directed to various organelle membranes, including the endoplasmic reticulum (ER) membrane, the mitochondrial outer membrane (OM), and peroxisomal membranes (Chio et al., 2017). The guided-entry of TA proteins (GET) pathway in yeast and the transmembrane recognition complex (TRC) pathway in mammals are the major targeting routes for TA proteins to the ER membrane (Schuldiner et al., 2008; Stefanovic and Hegde, 2007). In the GET pathway, the Sgt2 chaperone captures newly synthesized TA proteins from the ribosome and transfers them to the Get3 ATPase, with the aid of the Get4-Get5 complex. The Get3 loaded with TA proteins is directed to the Get1-Get2 complex on the ER membrane, where it releases the TA proteins. The Get1-Get2 complex then facilitates the insertion of the TA proteins into the ER membrane. The components of the GET pathway are well conserved in the TRC pathway, except for Bag6 (Hessa et al., 2011). In addition to the GET/TRC pathway, the ER membrane complex (EMC) pathway and the signal recognition particle-independent (SND) pathway may also be involved in the ER targeting of some TA proteins (Aviram et al., 2016; Guna et al., 2018).

Pex15 is a peroxisomal TA protein in yeast (Elgersma et al., 1997). Two possible peroxisomal targeting pathways for Pex15 have been suggested, as follows. One pathway involves direct targeting mediated by Pex19 and Pex3 (Halbach et al., 2006). Pex19 is a cytosolic chaperone that binds to newly synthesized peroxisomal membrane proteins and associates with the docking receptor, Pex3, on the peroxisomal membrane (Fang et al., 2004). Pex19 recognizes the membrane peroxisome targeting signal (mPTS) of peroxisomal membrane proteins (Sacksteder et al., 2000; Jones et al., 2004). The C-terminal globular domain of Pex19 interacts with the mPTS, which consists of short helical motifs with basic and/or hydrophobic residues (Rottensteiner et al., 2004; Schueller et al., 2010). The mPTS of Pex15 was suggested to be encrypted in its C-terminal region containing the TM segment (Halbach et al., 2006). The second pathway involves the initial targeting of Pex15 to the ER *via* the GET pathway, followed by vesicular transport to peroxisomes (Schuldiner et al., 2008). In this ER-mediated pathway, Pex19 is considered to function as a sorting receptor to generate pre-peroxisomal vesicles (Lam et al., 2010; van der Zand et al., 2010). Although the direct and indirect peroxisomal targeting pathways for Pex15 may not be mutually exclusive, it remains unclear whether Pex15 primarily utilizes one of the two pathways. In contrast, PEX26, the mammalian homolog of Pex15, is directly targeted to peroxisomes in a PEX19- and PEX3-dependent manner, but independently of the TRC pathway (Yagita et al., 2013). Biochemical analyses have shown that the N-terminal conserved amphipathic segments and the farnesylation at a C-terminal CaaX motif in PEX19 are important for its direct interaction with its substrate, PEX26 (Chen et al., 2014; Emmanouilidis et al., 2017; Oh et al., 2024).

TA proteins targeted to the ER or peroxisomes can be mislocalized to the mitochondrial OM by genetic mutations in the GET pathway or the Pex19-Pex3-dependent pathway (Schuldiner et al., 2008; Jonicas et al., 2009; Nuebel et al., 2021). Yeast Msp1 and its mammalian homolog ATAD1 are conserved AAA-ATPases localized to the mitochondrial OM and peroxisomes. Msp1/ATAD1 removes such mistargeted TA proteins from the OM (Nakai et al., 1993, Chen et al., 2014; Okreglak and Walter, 2014). Viral TA proteins, endogenous OM proteins involved in mitochondrial fission and apoptosis, and precursor proteins accumulated on the mitochondrial OM or clogged in the translocase of the mitochondrial OM (TOM) complex have also been reported as substrates for Msp1/ATAD1 (Weidberg and Amon, 2018; Winter et al., 2022; Zhou et al., 2023; He et al., 2023; Kim et al., 2024). Cryo-electron microscopy analyses revealed that Msp1/ATAD1 functions as an ATP-dependent membrane dislocase that extracts mistargeted proteins from the membrane by forming a spiral-shaped, homo-hexameric structure (Wohlever et al., 2017; Wang et al., 2020, Wang et al., 2022). Pex15Δ30, a mutant of Pex15 lacking the C-terminal 30 residues following the TM segment, which binds to Pex19, serves as a model substrate for Msp1-dependent degradation (Okreglak and Walter, 2014). How Msp1 recognizes mislocalized TA proteins is a key issue. It has been suggested that Msp1 recognizes a hydrophobic patch near the TM segment of Pex15Δ30 on the mitochondrial OM (Li et al., 2019), while endogenous Pex15 on the peroxisomal membrane may evade recognition by peroxisomal Msp1 as it binds to Pex3 on the membrane (Weir et al., 2017). In addition, Msp1 is unable to recognize artificially dimerized Pex15Δ30 (Dederer et al., 2019). These findings suggest that Msp1 recognizes orphaned TA proteins that lack a binding partner as mislocalized proteins (Wang and Walter, 2020).

Our previous studies have shown that Msp1 extracts mistargeted TA proteins such as Pex15Δ30 and authentic ER-TA proteins from the mitochondrial OM and then transfers them to the ER *via* the GET pathway (Matsumoto et al., 2019; Matsumoto et al., 2022). Once in the ER, Pex15Δ30 is ubiquitinated by the ER-localized E3 ubiquitin ligase Doa10 and subsequently degraded by the proteasome (Matsumoto et al., 2019). In contrast, authentic TA proteins like Frt1 and Gos1 escape degradation, and instead are transported to their respective destinations in the ER and Golgi (Matsumoto et al., 2019; Matsumoto et al., 2022). We therefore proposed that Msp1 is not only involved in the degradation of mistargeted proteins, but also acts as a proofreading factor that corrects the mislocalization of TA proteins by giving them a second chance to reach their proper destinations (Matsumoto and Endo, 2023).

In this study, we addressed the question of how Pex15 is targeted to peroxisomes and found that Pex15 was mistargeted to mitochondria in yeast cells lacking *PEX19* or *PEX3*, suggesting the crucial roles of Pex19 and Pex3 in Pex15 targeting to peroxisomes. Next, we found that Msp1 and the GET pathway facilitate the transfer of mistargeted Pex15 from the mitochondrial OM to the ER for degradation, in the absence of Pex19 and Pex3. We then constructed yeast strains with an auxin-inducible degron system that facilitates the rapid degradation of Pex19 and Pex3. Upon the addition of auxin, the peroxisomal targeting of newly synthesized Pex15 was significantly inhibited due to the rapid depletion of Pex19 and Pex3. However, after replenishment of Pex19, the mislocalized Pex15 moved from the mitochondrial OM to peroxisomes in an Msp1-dependent manner. Interestingly, we found that endogenous Pex15 on peroxisomes was also extracted from the membrane by peroxisomal Msp1 when Pex19 or Pex3 was rapidly depleted, suggesting that peroxisomal Msp1 constantly extracts Pex15 from the membrane, but Pex15 can return to peroxisomes *via* the Pex19-Pex3-dependent pathway. Collectively, our findings provide new insights into the peroxisomal localization of Pex15, emphasizing the coordinated roles of dual-localized Msp1 and the Pex19-Pex3-dependent pathway.

## Results

### Mislocalized Pex15 is destabilized in *pex19*- and *pex3*-deletion yeast cells

To analyze the localization of the authentic peroxisomal TA protein Pex15 at endogenous expression levels, we inserted the gene fragment encoding mNeonGreen (mNG), a bright monomeric yellow-green fluorescent protein, into the *PEX15* gene for its N-terminal tagging by using a CRISPR-Cas9 system (Okada et al., 2021). We confirmed that mNG-tagged Pex15 (mNG-Pex15) co-localized with the peroxisomal matrix marker, mCherry-PTS1, in wild-type (WT) cells (Fig. 1A). We then found that mutations in *MSP1* and the GET pathway did not affect the peroxisomal localization of mNG-Pex15 (Fig. 1A). Next, to facilitate the detection of endogenous Pex15 by immunoblotting, we inserted a gene fragment encoding the 3xFLAG-tag into the *PEX15* gene in the same manner as the mNG-tag insertion. We performed cycloheximide (CHX) chase experiments to analyze its stability and found that 3xFLAG-Pex15 is highly stable, not only in WT cells, but also in *msp1*11, *get3*11 and *get1*11*get2*11 cells (Fig. 1B).

**Figure 1.**
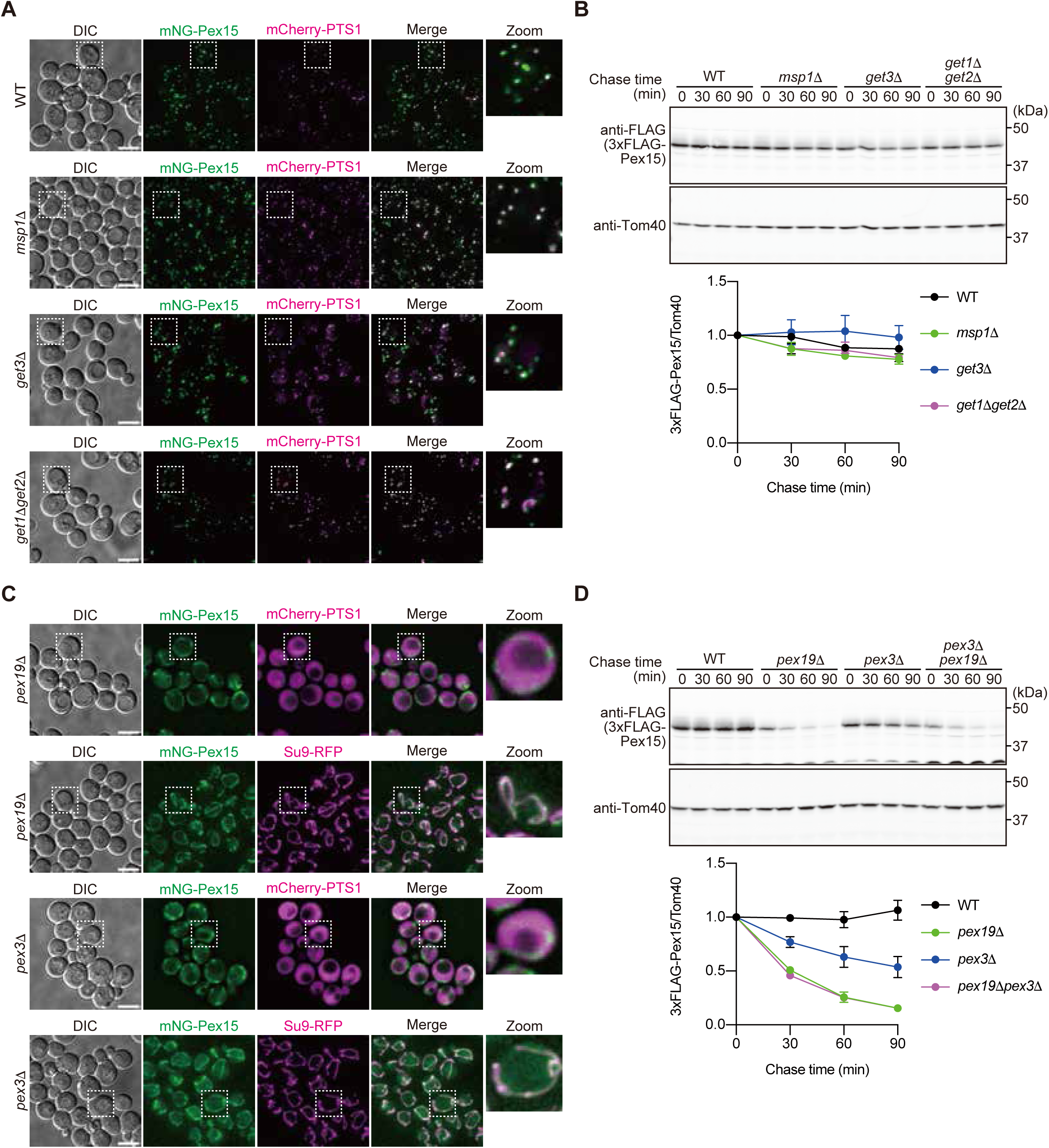
Deletions of *pex19* and *pex3* cause Pex15 mistargeting to mitochondria and its destabilization. **(A)** WT, *msp1*11, *get3*11 and *get1*11*get2*11 cells expressing mNG-Pex15 under the control of its own promoter from the chromosome, were grown in SCD medium at 30°C and imaged by fluorescence microscopy. Peroxisomes were labeled with mCherry-PTS1. Maximum projection images of Z-stacks are shown. Scale bar, 5 μm. DIC, differential interference contrast microscopy. **(B)** WT, *msp1*11, *get3*11 and *get1*11*get2*11 cells expressing 3xFLAG-Pex15 under the control of its own promoter from the chromosome, were grown in YPD medium at 30°C. At the indicated times after the addition of 100 µg/ml CHX, cell extracts were prepared and proteins were analyzed by SDS-PAGE and immunoblotting with the indicated antibodies (left). The normalized relative amounts of 3xFLAG-Pex15 were plotted against the chase time. Values are the mean ± S.D. from three independent experiments. **(C)** *pex19*11 and *pex3*11 cells expressing mNG-Pex15 under the control of its own promoter from the chromosome, were grown in SCD medium at 30°C and imaged by fluorescence microscopy. Peroxisomes and mitochondria were labeled with mCherry-PTS1 and Su9-RFP, respectively. Single-plane for peroxisomes and maximum projection for mitochondria images of Z-stacks are shown. Scale bar, 5 μm. DIC, differential interference contrast microscopy. **(D)** WT, *pex19*11, *pex3*11 and *pex19*11*pex3*11 cells expressing 3xFLAG-Pex15 under the control of its own promoter from the chromosome, were grown in YPD medium at 30°C. At the indicated times after the addition of 100 µg/ml CHX, cell extracts were prepared and proteins were analyzed by SDS-PAGE and immunoblotting with the indicated antibodies (left). The normalized relative amounts of 3xFLAG-Pex15 were plotted against the chase time. Values are the mean ± S.D. from three independent experiments.

Next, we analyzed the localization and stability of Pex15 in *PEX19-* and *PEX3*-deficient yeast cells. The peroxisomal matrix marker, mCherry-PTS1, exhibited a cytosolic distribution because of the absence of peroxisomal membranes in *pex19*11 and *pex3*11 cells (Fig. 1C). In contrast, mNG-Pex15 co-localized with the mitochondrial matrix marker, Su9-RFP, in these mutant cells (Fig. 1C). A CHX chase analysis showed that 3xFLAG-Pex15 was destabilized in *pex19*11 and *pex3*11 cells compared to WT (Fig. 1D), and its half-lives in *pex19*11 and *pex19*11*pex3*11 cells were shorter than that in *pex3*11 cells (Fig. 1D). In WT cells, the N-terminally mNG-tagged Pex19 (mNG-Pex19) was localized to both the peroxisomes and cytosol in WT cells, whereas in *pex3*11 cells, mNG-Pex19 was localized exclusively in the cytosol (Fig. S1). These results suggest that Pex19 protects mislocalized Pex15 from degradation in the cytosol in *pex3*11 cells.

### Degradation pathway for mislocalized Pex15

We next searched for factors involved in the degradation of Pex15 mislocalized to mitochondria by the absence of Pex19. We first found that the degradation of 3xFLAG-Pex15 in *pex19*11 cells was inhibited by the addition of MG132, a proteasome inhibitor, in the absence of *PDR5*, encoding a multidrug exporter (Fig. S2), suggesting that mislocalized Pex15 is degraded by the proteasome like Pex151130 (Matsumoto et al., 2019). We next asked if Msp1 and the GET pathway are involved in the degradation of mislocalized Pex15 in *pex19*11 cells, and found that 3xFLAG-Pex15 was highly stabilized in *pex19*11*msp1*11 and *pex19*11*get3*11 cells (Fig. 2A). The mitochondrial signals of mNG-Pex15 were more pronounced in *pex19*11*msp1*11 cells than in *pex19*11 cells (Figs. 1C and 2B), likely due to the suppression of Msp1-mediated extraction of mNG-Pex15 from the mitochondrial OM. In contrast, in *pex19*11*get3*11 cells, the mNG-Pex15 fluorescent signals were observed at mitochondria as well as cytosolic puncta (Fig. 2B). These cytosolic puncta were partially co-localized with Hsp104-mCherry (Fig. 2C), indicating that the Pex15 mistargeted to the mitochondrial OM is extracted by Msp1, but the extracted Pex15 forms protein aggregates in the cytosol when the GET pathway is blocked.

**Figure 2.**
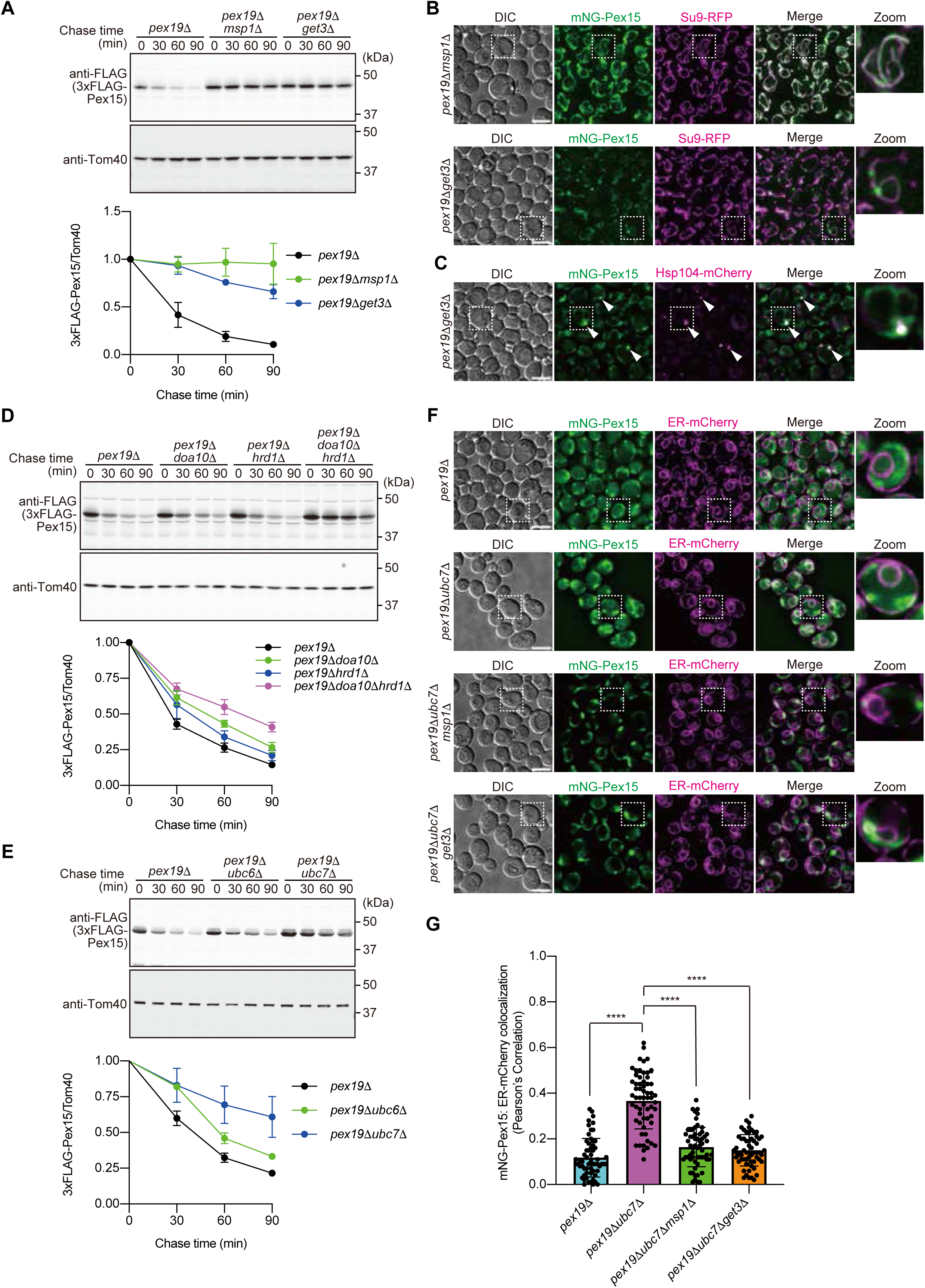
Mistargeted Pex15 is transferred to the ER *via* Msp1 and the GET pathway and then degraded by the ERAD system. **(A)** *pex19*11, *pex19*11*msp1*11 and *pex19*11*get3*11 cells expressing 3xFLAG-Pex15 under the control of its own promoter from the chromosome, were grown in YPD medium at 30°C. CHX-chase experiments were performed as in Fig. 1B. **(B)** *pex19*11*msp1*11 and *pex19*11*get3*11 cells expressing mNG-Pex15 under the control of its own promoter from the chromosome, were grown in SCD medium at 30°C and imaged by fluorescence microscopy as in Fig. 1A. **(C)** *pex19*11*get3*11 cells expressing mNG-Pex15 and Hsp104-mCherry under the control of their own promoters from the chromosomes, were grown in SCD medium at 30°C and imaged by fluorescence microscopy as in Fig. 1A. **(D)** *pex19*11, *pex19*11*doa10*11, *pex19*11*hrd1*11 and *pex19*11*doa10*11*hrd1*11 cells expressing 3xFLAG-Pex15 under the control of its own promoter from the chromosome, were grown in YPD medium at 30°C. CHX-chase experiments were performed as in Fig. 1B. **(E)** *pex19*11, *pex19*11*ubc6*11 and *pex19*11*ubc7*11 cells expressing 3xFLAG-Pex15 under the control of its own promoter from the chromosome, were grown in YPD medium at 30°C. CHX-chase experiments were performed as in Fig. 1B. **(F)** *pex19*11, *pex19*11*ubc7*11, *pex19*11*ubc7*11*msp1*11 and *pex19*11*ubc7*11*get3*11 cells expressing mNG-Pex15 under the control of its own chromosomal promoter from the chromosome, were grown in SCD medium at 30°C and imaged by fluorescence microscopy. The ER was labeled with BipN-mCherry-HDEL (ER-mCherry). Single-plane images are shown. Scale bar, 5 μm. DIC, differential interference contrast microscopy. **(G)** Colocalization of mNG-Pex15 with the ER was quantified by using Pearson’s correlation coefficient between the mNG and ER (mCherry) signals. Values are the mean ± S.D. (*n*=60) from three independent replicates; *n* represents the number of cells. ****, P < 0.0001 compared with *pex19*11*ubc7*11 cells by one-way ANOVA with Dunnett’s multiple comparison test.

As in the case of Pex151130, we examined if Doa10 is involved in the degradation of Pex15 transferred to the ER. Contrary to our expectations, we found that the degradation of 3xFLAG-Pex15 was not strongly inhibited in *pex19*11*doa10*11 cells (Fig. 2D). Next, we asked if Hrd1, another E3 ligase in the ERAD pathway, is involved in the degradation of mislocalized Pex15, and found that the degradation rate of 3xFLAG-Pex15 was slightly delayed in *pex19*11*hrd1*11 cells compared to *pex19*11 cells (Fig. 2D). To explore the possibility of Doa10 and Hrd1 serving as overlapping E3 ligases for Pex15 in the ER, we further assessed the stability of 3xFLAG-Pex15 in *pex19*11*doa10*11*hrd1*11 cells. The degradation of 3xFLAG-Pex15 was more strongly impaired in *pex19*11*doa10*11*hrd1*11 cells than in *pex19*11 cells lacking either Doa10 or Hrd1 alone (Fig. 2D). Consistent with this result, the degradation of 3xFLAG-Pex15 in *pex19*11 cells was more strongly inhibited when Ubc7, the shared E2 enzyme of Doa10 and Hrd1, was deleted, as compared to the deletion of the gene encoding Ubc6, the E2 enzyme specific to Doa10 (Fig. 2E). In support of these results, a portion of the mNG-Pex15 signal was co-localized with the ER in *pex19*11*ubc7*11 cells, but not in *pex19*11 cells (Figs. 2F, G). In addition, the ER-localization of mNG-Pex15 in *pex19*11*ubc7*11 cells was suppressed by the combined mutation with either *MSP1* or *GET3* (Figs. 2F, G). Taken together, these analyses indicated that Msp1 and the GET pathway mediate the transfer of mislocalized Pex15 from mitochondria to the ER for degradation by the system.

### Pex19 and Pex3 are involved in the peroxisomal targeting of Pex15

Our results suggested that Pex15 utilizes the Pex19-Pex3-dependent pathway for targeting to peroxisomes. To demonstrate the direct involvement of Pex19 and Pex3 in the peroxisomal targeting of Pex15, the effect of regulating the Pex19 and Pex3 expression levels on the peroxisomal targeting of Pex15 must be examined while maintaining the integrity of peroxisomal membranes. We previously developed a method, termed the ‘Turn-ON/OFF system’, to analyze the Msp1-dependent transfer of mistargeted TA proteins from mitochondria to the ER (Matsumoto et al., 2022). In this study, we depleted Pex19 or Pex3 using rapid degradation *via* the auxin-inducible degron (AID) system, which facilitates the acute loss of the target protein specified by the AID tag upon the addition of auxin, indole-3-acetic acid (IAA) (Nishimura et al., 2009) (Fig. 3A and B). First, we inserted the minimal AID-tag (AID*) just downstream of the start codon in the *PEX19* genes, using the CRISPR-Cas9 system in OsTIR1-expressing yeast cells. The functionality of the N-terminally AID-tagged Pex19 (AID*-Pex19) was not affected, since peroxisomal membrane markers (Pex11-mCherry) remained visible as fluorescent dots in the absence of IAA (Fig. 3D). We confirmed that AID*-Pex19 is rapidly depleted (‘Turn-OFF’) within 30 minutes by IAA treatment (Fig. 3C). Next, we induced 3xFLAG-tagged-mNG-Pex15 (‘Turn-ON’) expression under the control of the cyanamide-inducible *DDI2* promoter in AID*-Pex19-expressing yeast cells, and analyzed its localization in the presence (-IAA) and absence (+IAA) of AID*-Pex19. The results showed that cyanamide-induced 3xFLAG-mNG-Pex15 co-localized with peroxisomal markers in the presence of AID*-Pex19 (-IAA), but this co-localization was significantly reduced in the absence of AID*-Pex19 (+IAA) (Figs. 3D and E). We then deleted the *MSP1* gene in the AID*-Pex19-expressing yeast strain. In this strain, cyanamide-induced 3xFLAG-mNG-Pex15 is localized to peroxisomes in the presence of AID*-Pex19 (-IAA), but is accumulated in mitochondria in the absence (+IAA) of AID*-Pex19 (Figs. S3 A-D). We next constructed the AID system for Pex3 and confirmed the rapid IAA-dependent depletion of Pex3-AID*-V5 (Fig. 3F). Furthermore, the peroxisomal localization of cyanamide-induced 3xFLAG-mNG-Pex15 was inhibited in the absence of Pex3-AID*-V5 (+IAA) (Figs. 3G and H). In contrast, using the AID system of Get3 (Get3-AID*-9xMyc), we found that Pex15 was localized to peroxisomes even in the absence of Get3-AID*-9xMyc (Figs. S3 E and F). These results indicate that the targeting of Pex15 to peroxisomes is dependent on Pex19 and Pex3, but not on the GET pathway, suggesting that Pex15 is transported to peroxisomes *via* a direct route involving Pex19 and Pex3.

**Figure 3.**
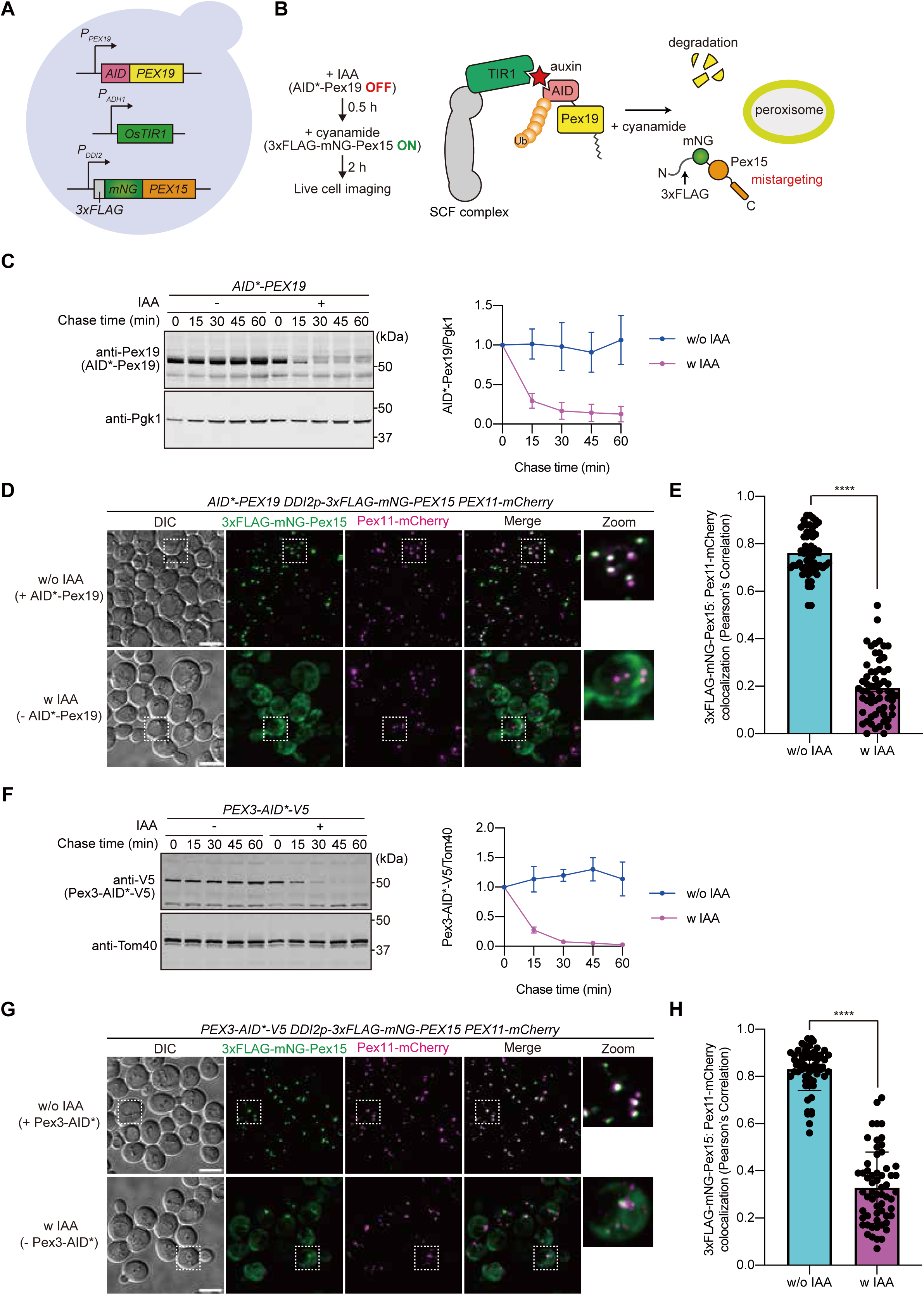
Newly synthesized Pex15 is targeted to peroxisomes in a Pex19- and Pex3-dependent manner. **(A)** A schematic representation of the acute depletion of Pex19 using the AID system. N-terminally AID*-tagged Pex19 was expressed under the control of its own promoter from the chromosome. 3xFLAG-mNG-Pex15 was expressed from the cyanamide-inducible *DDI2* promoter and OsTIR1 was expressed from the *ADH1* promoter. **(B)** AID*-Pex19 was depleted by a treatment with 1 mM IAA for 30 min at 30°C, followed by the induction of 3xFLAG-mNG-Pex15 with 5 mM cyanamide at 30°C for 2 hours. The localization of 3xFLAG-mNG-Pex15 was then analyzed using fluorescence microscopy. **(C)** Yeast cells in Fig. 3A were grown in SCD medium at 30°C. Cell extracts were prepared at the indicated times in the presence (-IAA) or absence (+IAA) of AID*-Pex19. Proteins were analyzed by SDS-PAGE and immunoblotting with the indicated antibodies. The relative amounts of AID*-Pex19 (upper panel) and Pgk1 (lower panel) were plotted against the chase time. Values are the mean ± S.D. from three independent experiments. (**D**) The localization of 3xFLAG-mNG-Pex15 was imaged by fluorescence microscopy in the presence (-IAA, upper panel) or absence (+IAA, lower panel) of AID*-Pex19. Maximum projection images of Z-stacks are shown. Peroxisomes were labeled with Pex11-mCherry. Scale bar, 5 μm. DIC, differential interference contrast microscopy. (**E**) Colocalization of 3xFLAG-mNG-Pex15 with peroxisomes was quantified using Pearson’s correlation coefficient between the mNG and mCherry signals. Values are the mean ± S.D. (-IAA; *n*=60, +IAA; *n*=60) from three independent replicates; *n* represents the number of cells. ****, P < 0.0001 compared with IAA untreated (-IAA) cells by two-tailed paired *t* test. **(F)** Yeast cells expressing Pex3-AID*-V5 under the control of its own promoter from the chromosome were grown in SCD medium at 30°C. Extracts were prepared at the indicated times from cells cultured in the presence (+IAA) or absence (-IAA) of Pex3-AID*-V5, and proteins were analyzed by SDS-PAGE and immunoblotting with the indicated antibodies. The relative amounts of Pex3-AID*-V5 (upper panel) and Tom40 (lower panel) were plotted against the chase time. Values are the mean ± S.D. from three independent experiments. **(G)** The localization of 3xFLAG-mNG-Pex15 in the absence (-IAA, upper panel) or presence (+IAA, lower panel) of Pex3-AID*-V5 was imaged by fluorescence microscopy as in (B). Maximum projection images of Z-stacks are shown. Peroxisomes were labeled with Pex11-mCherry. Scale bar, 5 μm. DIC, differential interference contrast microscopy. (**H**) Colocalization of 3xFLAG-mNG-Pex15 with peroxisomes was quantified using Pearson’s correlation coefficient between the mNG and mCherry signals, as in (E). Values are the mean ± S.D. (-IAA; *n*=60, +IAA; *n*=60) from three independent replicates; *n* represents the number of cells. ****, P < 0.0001 compared with IAA untreated (-IAA) cells by two-tailed paired *t* test.

### Mislocalized Pex15 can move from mitochondria to peroxisomes in an Msp1- and Pex19-dependent manner

We next asked if Pex15, which is extracted from the mitochondrial OM by Msp1, can be transferred to peroxisomes in a Pex19- and Pex3-dependent manner. We thus utilized a yeast strain that can rapidly degrade Pex19 by the AID system (Fig. 4A). Due to the dual localization of Msp1 to both the mitochondrial OM and peroxisomal membrane, it is possible that Pex15, once returned to peroxisomes from mitochondria, could be extracted again from the peroxisomal membrane by Msp1 on peroxisomes. To minimize this possibility, we deleted the endogenous *MSP1* gene and expressed a mitochondrially targeted Msp1 variant (Tom70TM-Msp1), in which the N-terminal TM segment of Msp1 was replaced with that of Tom70 under the control of the β-Estradiol-inducible *Z4EV* promoter (Fig. 4A).

**Figure 4.**
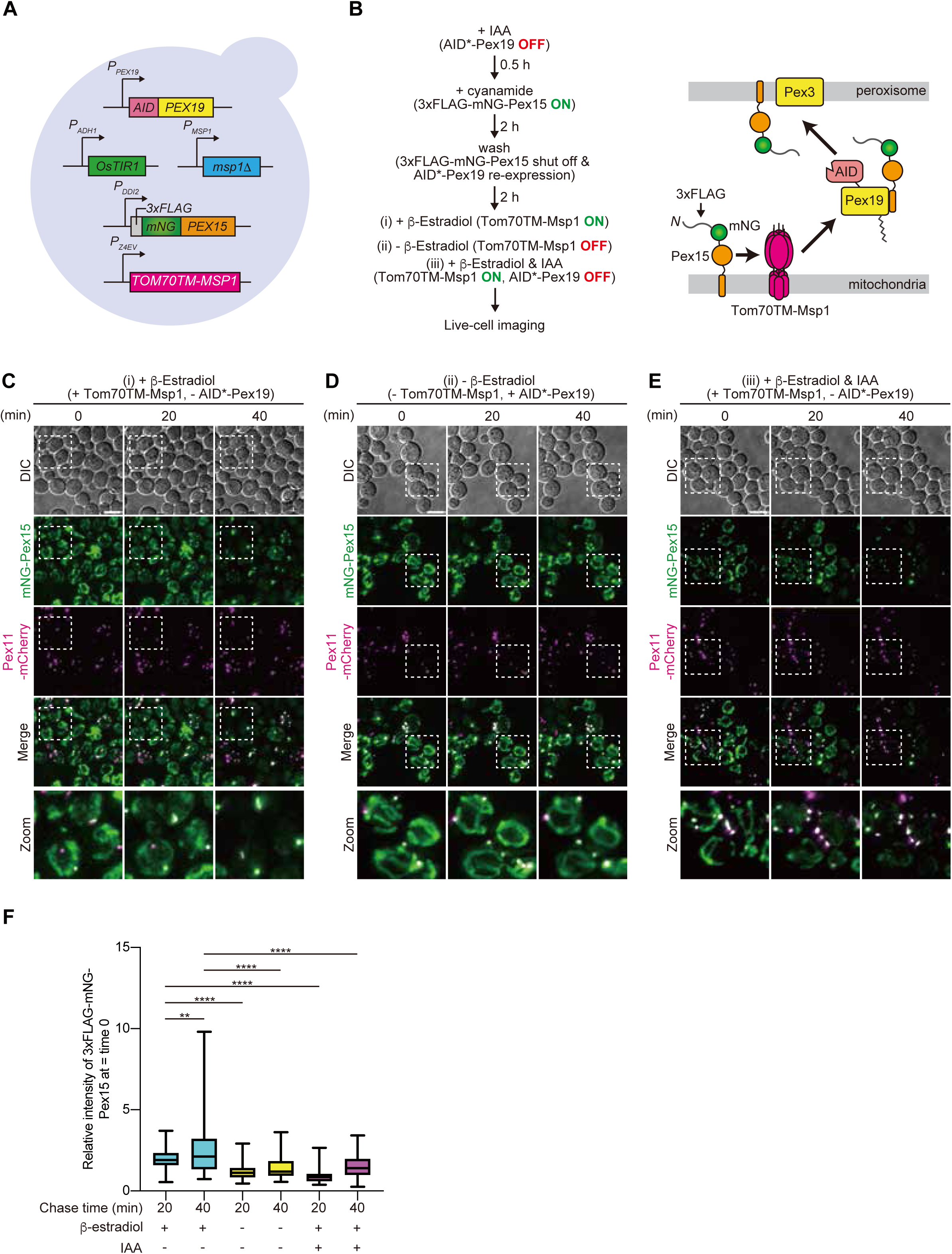
Msp1- and Pex19-dependent transfer of mistargeted Pex15 from mitochondria to peroxisomes. **(A)** A schematic representation of the live-cell imaging. As in Fig. 3A, AID*-Pex19 and OsTIR1 were expressed under the control of its own promoter and the *ADH1* promoter, respectively. 3xFLAG-mNG-Pex15 was expressed under the control of the cyanamide-inducible *DDI2* promoter. The endogenous *MSP1* gene was deleted to block the extraction of Pex15 from peroxisomes. Instead, a mitochondria-targeted Msp1 variant (Tom70TM-Msp1) was expressed under the control of the β-Estradiol-inducible *Z4EV* promoter. **(B)** Yeast cells in Fig. 4A were grown in SCD medium at 30°C, and AID*-Pex19 was depleted by a treatment with 1 mM IAA for 30 min at 30°C. 3xFLAG-mNG-Pex15 was then induced with 5 mM cyanamide for 2 h at 30°C. The cells were washed with fresh SCD medium to shut off the 3xFLAG-mNG-Pex15 expression and allow the re-expression of AID*-Pex19 by growth in SCD medium for 2 h at 30°C. Time-lapse imaging was recorded after the addition of 1 μM β-Estradiol to induce the expression of Tom70TM-Msp1. **(C)** Yeast cells in (A) were grown in SCD at 30°C, and treated with 1 μM β-Estradiol after the expression of 3xFLAG-mNG-Pex15 was shut off. Time-lapse images were then obtained at 20-minute intervals. Maximum projection images of Z-stacks are shown. Peroxisomes were labeled with Pex11-mCherry. Scale bar, 5 μm. **(D)** Time-lapse microscopy images were obtained at the indicated times (min) for the cells in (A) without IAA and β-Estradiol treatment. Maximum projection images of Z-stacks are shown. Peroxisomes were labeled with Pex11-mCherry. Scale bar, 5 μm. **(E)** Yeast cells in A were grown in SCD at 30°C, and treated with 1 mM IAA and 1 μM β-Estradiol after the expression of 3xFLAG-mNG-Pex15 was shut off. Maximum projection images of Z-stacks are shown. Peroxisomes were labeled with Pex11-mCherry. Scale bar, 5 μm. **(F)** For the microscopic images obtained in (C-E), the fluorescence intensity of 3xFLAG-mNG-Pex15 on peroxisomes (labeled with Pex11-mCherry) was quantified at each time point. A single yeast cell was selected as a region of interest (ROI), and the intensity was shown relative to Time 0 in cells extracted as ROIs. Values are the mean ± S.D. (+β-Estradiol; *n*=60, -β-Estradiol; *n*=60, +IAA, β-Estradiol; *n*=60) from three independent replicates; *n* represents the number of cells. ****, P < 0.0001; **, P = 0.0475 compared with the results of time-lapse imaging with β-Estradiol treatment by one-way ANOVA with Dunnett’s multiple comparison test.

We first treated the cells with IAA to rapidly remove AID*-Pex19 (‘Turn-OFF’), and then induced 3xFLAG-mNG-Pex15 (‘Turn-ON’) through cyanamide addition (Fig. 4B). Next, both IAA and cyanamide were removed from the medium to promote Pex19 re-expression and to shut off 3xFLAG-mNG-Pex15 expression, respectively (Fig. 4B). After a 2 h incubation to attenuate Pex15 transcription and re-express AID*-Pex19, we induced Tom70TM-Msp1 expression (‘Turn-ON’) by the addition of β-Estradiol (Fig. 4B). In our previous study, we confirmed that the mRNA level of cyanamide-induced Pex15Δ30 decreased to about 5% of that immediately after induction during the 2 h incubation, as measured by quantitative PCR (Matsumoto et al., 2022). To analyze the localization changes of 3xFLAG-mNG-Pex15 from mitochondria to peroxisomes, we performed live-cell imaging using three different conditions (Fig. 4B). We observed a gradual increase in the 3xFLAG-mNG-Pex15 signal (green) on peroxisomes (magenta) upon the induction of Tom70TM-Msp1 expression by the addition of β-Estradiol (+β-Estradiol; Figs. 4C, F). In contrast, without Tom70TM-Msp1 expression (-β-Estradiol; Figs. 4D, F), the 3xFLAG-mNG-Pex15 signals remained on mitochondria. These results suggest that the Pex15 on mitochondria is extracted by the mitochondrially localized Msp1 and transferred to peroxisomes.

We next investigated the role of Pex19 in the Msp1-dependent transfer of 3xFLAG-mNG-Pex15 from mitochondria to peroxisomes. When Msp1 was expressed (+β-Estradiol) and Pex19 was depleted (+IAA), the 3xFLAG-mNG-Pex15 signal decreased on mitochondria but did not increase on peroxisomes (Figs. 4E, F). These results indicated that 3xFLAG-mNG-Pex15 was extracted from the OM by Msp1 but did not reach peroxisomes due to the absence of Pex19, suggesting that Pex19 is important for the transfer of mislocalized Pex15 from mitochondria to peroxisomes. We then tracked the protein levels of Pex15 mislocalized to the mitochondrial OM after Msp1 expression (+β-Estradiol), in the presence and absence of Get3 (Fig. S4). The decrease in the Pex15 protein level was less pronounced by the loss of Get3, despite the restored expression of Pex19. These findings suggest that the Pex15 extracted from the OM by Msp1 is not only transferred to peroxisomes in a Pex19-dependent manner, but also partially transported to the ER *via* the GET pathway for degradation.

### Msp1-dependent turnover of ‘endogenous’ Pex15 on peroxisomes under Pex19-depleted conditions

Pex15 binding to Pex3 on peroxisomal membranes reportedly prevents recognition by Msp1 on peroxisomes (Weir et al., 2017). Consistently, our results demonstrated that the Pex15 on peroxisomes at endogenous expression levels is highly stable (Fig. 1A). Here, we showed that both Pex19 and Pex3 are involved in the peroxisomal targeting of newly synthesized Pex15 (Fig. 3) and the transfer of mistargeted Pex15 from mitochondria to peroxisomes (Fig. 4). Based on these observations, we hypothesized that Pex19 functions as a chaperone for Pex15 in the cytosol, even when it is extracted by Msp1 on peroxisomes, facilitating its return by passing it to Pex3 on peroxisomes. To test this possibility, we tagged ‘endogenous’ Pex15 with a 3xFLAG tag at its N-terminus in AID*-Pex19-expressing yeast strains and examined the stability of peroxisome-localized Pex15 in the presence (-IAA) or absence (+IAA) of AID*-Pex19 by CHX chase experiments. We found that 3xFLAG-Pex15 was partially degraded in the presence of AID*-Pex19 (-IAA), but the degradation of 3xFLAG-Pex15 was significantly accelerated under AID*-Pex19-depleted conditions (Fig. 5A). Moreover, the deletion of *MSP1* suppressed the destabilization of 3xFLAG-Pex15 in the absence of AID*-Pex19 (+IAA) (Fig. 5A). To confirm that the degradation of endogenous Pex15 occurs on peroxisomes upon Pex19 depletion, we tagged Pex15 with mNG at its N-terminus in AID*-Pex19-expressing yeast cells and tracked its localization by a CHX chase and fluorescence microscopy in the absence (+IAA) of AID*-Pex19. The mNG-Pex15 signal (green) on peroxisomes (magenta) rapidly disappeared in Msp1-expressing cells, but remained on peroxisomes for a longer time in the absence of Msp1 (Fig. 5B).

**Figure 5.**
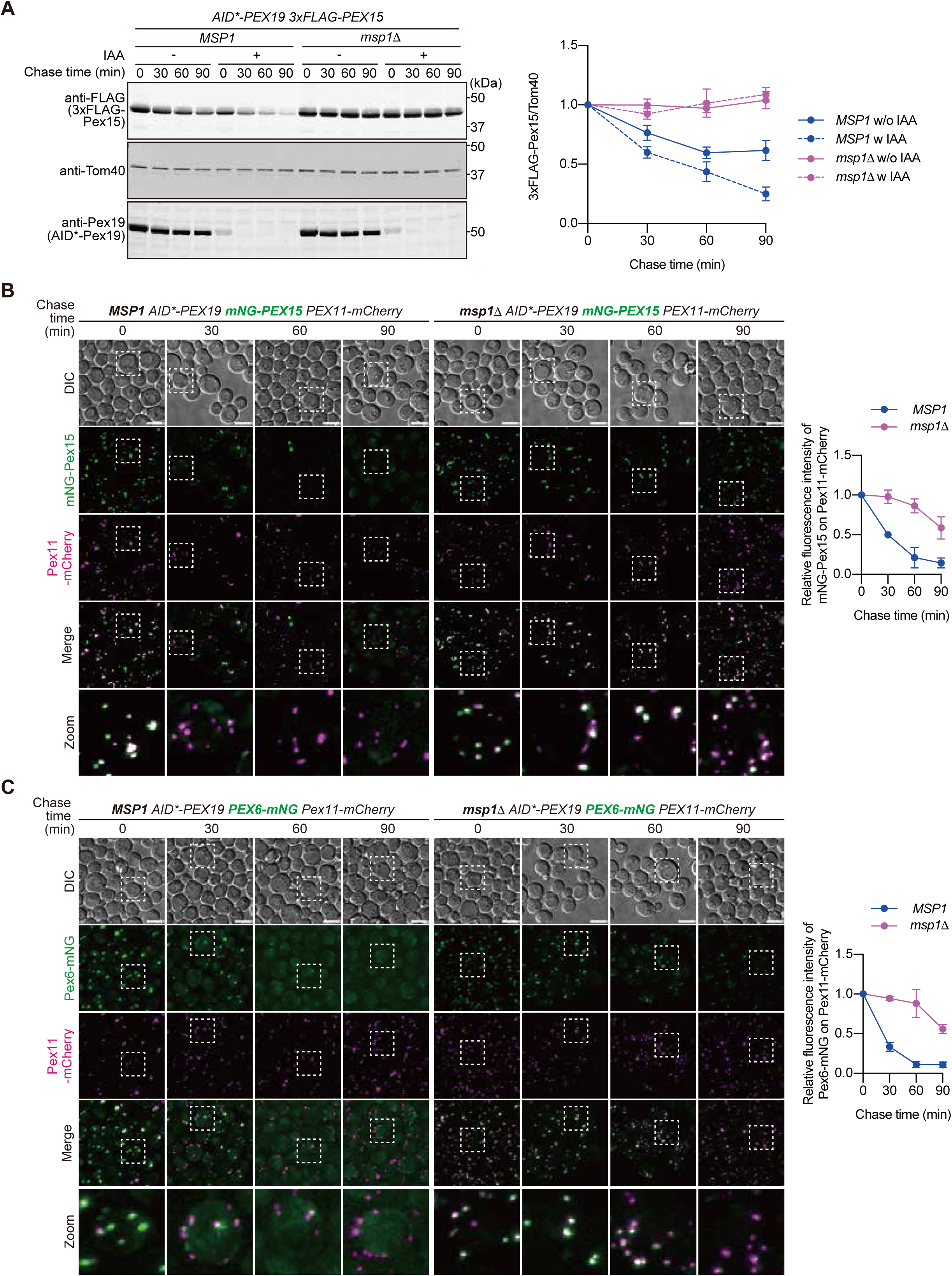
Peroxisomal Msp1 mediates endogenous Pex15 degradation under rapid Pex19 depletion conditions. **(A)** Wild-type (*MSP1*) and *msp1*11 cells expressing AID*-Pex19 and 3xFLAG-Pex15 under the control of their own promoters from the chromosomes were grown in YPD medium at 30°C, and then further incubated for 30 min after the addition of 1 mM IAA. Extracts were prepared at the indicated times after the addition of 100 µg/ml CHX, and proteins were analyzed by SDS-PAGE and immunoblotting with the indicated antibodies (left). The normalized relative amounts of 3xFLAG-Pex15 were plotted against the chase time. Values are the mean ± S.D. from three independent experiments. **(B)** Wild-type (*MSP1*) and *msp1*11 cells expressing AID*-Pex19 and mNG-Pex15 under the control of their own promoters from the chromosomes were grown in SCD medium at 30°C. The localization of mNG-Pex15 was observed by fluorescence microscopy at the indicated time points after the addition of 1 mM IAA and 100 µg/ml CHX. Maximum projection images of Z-stacks are shown. Peroxisomes were labeled with Pex11-mCherry. Scale bar, 5 μm. DIC, differential interference contrast microscopy. Background subtraction was first performed on the microscopic images, and followed by z-stack maximum intensity projection. The mNG-Pex15 fluorescence signal on peroxisomes was then quantified from more than 100 cells at each time point per experiment. The average intensity of the mNG-Pex15 fluorescence signal was calculated and plotted as a relative value, with the signal at time 0 set to 1. The graph shows the averages of data from two independent experiments. **(C)** Wild-type (*MSP1*) and *msp1*11 cells expressing AID*-Pex19 and Pex6-mNG under the control of their own promoters from the chromosomes were grown in SCD medium at 30°C. The localization of Pex6-mNG was observed by fluorescence microscopy at the indicated time points after the addition of 1 mM IAA and 100 µg/ml CHX. Maximum projection images of Z-stacks are shown. Peroxisomes were labeled with Pex11-mCherry. Scale bar, 5 μm. DIC, differential interference contrast microscopy. The quantification of the Pex6-mNG signal was performed as in (B), with values plotted relative to the signal at time 0 set to 1. The averages from two independent experiments are shown.

Next, we analyzed the stability and localization of endogenous Pex15 following the rapid depletion of Pex3, using a similar approach to that for Pex19. Unlike the results obtained with the rapid depletion of AID*-Pex19, the total protein levels of 3xFLAG-Pex15 remained unchanged in both the presence (-IAA) and absence (+IAA) of Pex3-AID*-V5 (Fig. S5A). However, the mNG-Pex15 signal (green) on peroxisomes (magenta) was rapidly lost upon Pex3-AID*-V5 depletion in the Msp1-expressing cells, whereas the mNG-Pex15 signal remained on peroxisomes in cells lacking *MSP1* (Fig. S5B). Finally, we explored the possibility that under conditions of AID*-Pex19 deletion (+IAA), Pex3 may become destabilized, leading to Pex15 becoming orphaned on peroxisomes. To test this, we examined the stability of Pex3 tagged with 3xFLAG in the presence (-IAA) and absence (+IAA) of AID*-Pex19, and found that Pex3-3xFLAG remained stable under both conditions (Fig. S5C). These analyses suggest that the Msp1 on peroxisomes continuously extracts the endogenous Pex15 from the peroxisomal membrane and releases it into the cytosol, but it is subsequently returned to peroxisomes through a Pex19- and Pex3-dependent mechanism.

### Msp1 prevents peroxisomal recruitment of the Pex1-Pex6 complex under Pex19-depleted conditions

Pex15 acts as a scaffold for the hetero-hexameric AAA-ATPase Pex1-Pex6 complex on the peroxisomal membrane (Rosenkranz et al., 2006). This Pex1-Pex6 complex is an essential factor for protein import into the peroxisomal matrix, functioning in the ATP-dependent export of ubiquitinated Pex5 back to the cytosol (Platta et al., 2024). The interaction between Pex15 and the Pex1-Pex6 complex is mediated by the direct association between the N-terminal domain of Pex6 and the cytosolic region of Pex15 (Birschmann et al., 2003; Ali et al., 2024). We next examined whether the Msp1 on peroxisomes dissociates the Pex1-Pex6 complex from the peroxisomal membrane, by extracting Pex15 when Pex19 is rapidly depleted. For this purpose, we tagged Pex6 with mNG at its C-terminus in AID*-Pex19-expressing yeast cells, and analyzed its localization during a CHX chase with the addition of IAA, as outlined in Fig. 5C. The Pex6-mNG signal (green) on peroxisomes (magenta) rapidly decreased in the Msp1-expressing cells, whereas in *msp1*11 cells, the Pex6-mNG signal remained relatively stable on peroxisomes (Fig. 5C). These results suggest that the peroxisomal exportomer complex, consisting of Pex1-Pex6-Pex15, is immediately disassembled by the action of Msp1 on peroxisomes in the absence of Pex19, which likely prevents protein import into the peroxisomal matrix.

### Peroxisomal Msp1 mediates the transfer of Pex15 from peroxisomes to mitochondria

We wondered whether the endogenous Pex15 could be mistargeted to non-peroxisomal organelles such as mitochondria if the Msp1 extraction on peroxisomes is predominant or the Msp1 is exclusively present on the peroxisomal membrane. To test this possibility, we expressed a peroxisome-targeted Msp1 variant, Pex22TM-Msp1, and a mitochondria-targeted variant, Tom70TM-Msp1, as a control under the endogenous *MSP1* promoter in *msp1*11 cells (Fig. 6A). We then analyzed the localization of mNG-Pex15 expressed from its own promoter, using fluorescence microscopy. No mitochondrial localization of mNG-Pex15 was observed in WT, *msp1*11, or *msp1*11 cells expressing Tom70TM-Msp1 (Figs. 6B, C). In contrast, the mitochondrial localization of mNG-Pex15 was significantly increased when Pex22TM-Msp1 was expressed in *msp1*11 cells (Figs. 6B, C). These results support the model that peroxisomal Msp1 facilitates the transfer of Pex15 from peroxisomes to mitochondria. Thus, the expression of Msp1 on peroxisomes may increase the risk of Pex15 mistargeting to mitochondria.

**Figure 6.**
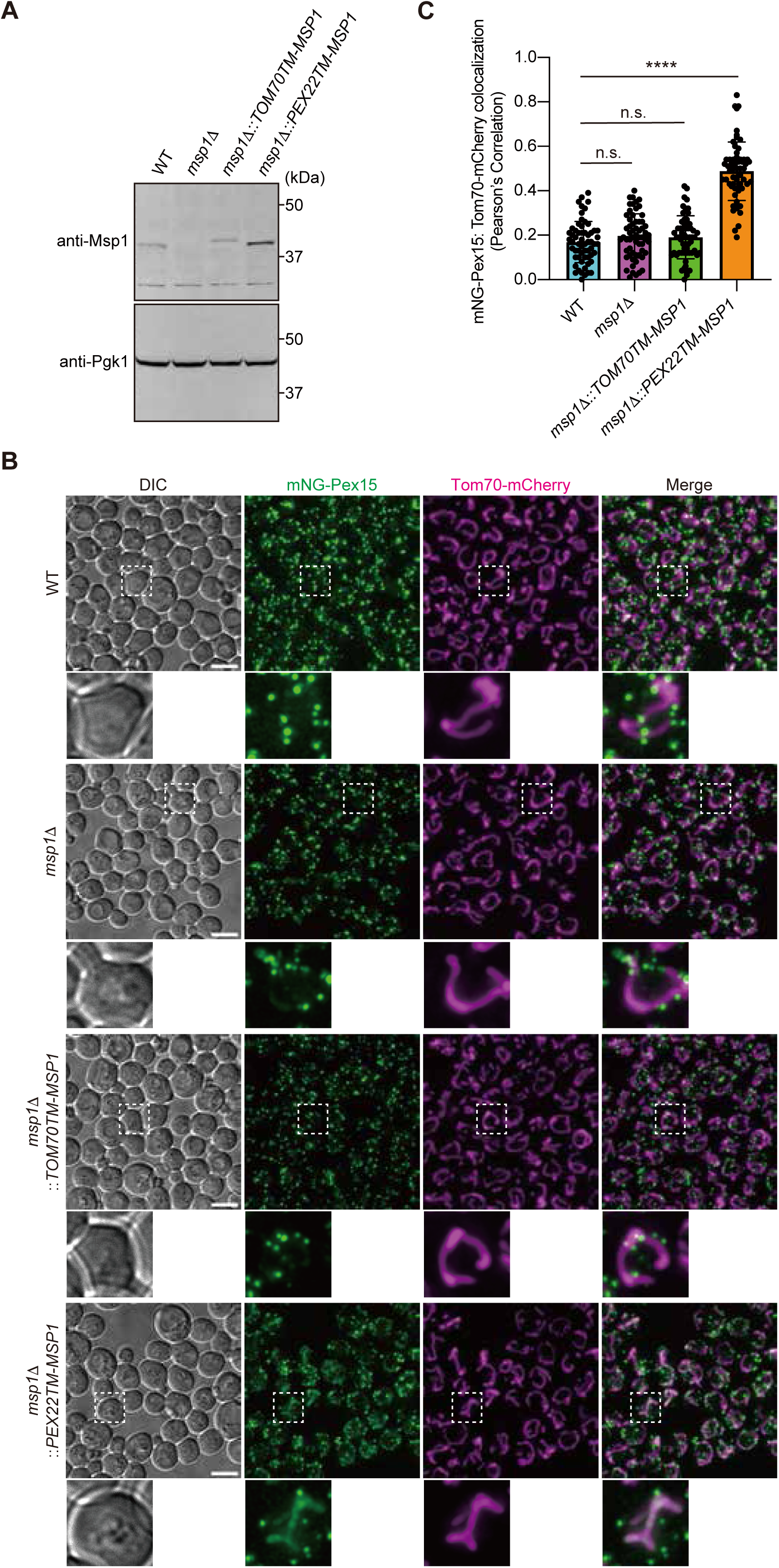
Peroxisomal Msp1 transfers endogenous Pex15 from peroxisomes to mitochondria. **(A)** Wild-type and *msp1*11 cells expressing mNG-Pex15 under the control of its own promoter from the chromosome were grown SCD medium at 30°C. In *msp1*11 cells, Tom70TM-Msp1 or a peroxisome-targeted Msp1 variant, Pex22TM-Msp1, was expressed under the control of its own promoter from the chromosome. Cell extracts were prepared, and proteins were analyzed by SDS-PAGE and immunoblotting with the indicated antibodies (left). **(B)** Yeast cells in (A) were grown in SCD medium at 30°C and imaged by fluorescence microscopy. Mitochondria were labeled with Tom70-mCherry. Maximum projection images of Z-stacks are shown. Scale bar, 5 μm. DIC, differential interference contrast microscopy. **(C)** Colocalization of mNG-Pex15 with mitochondria was quantified by using Pearson’s correlation coefficient between the mNG and mitochondria (mCherry) signals. Values are the mean ± S.D. (*n*=60) from three independent replicates; *n* represents the number of cells. ****, P < 0.0001 compared with wild-type by one-way ANOVA with Dunnett’s multiple comparison test.

## Discussion

We identified the targeting pathway for the peroxisomal TA protein, Pex15, and its degradation or transfer mechanisms when mislocalized to mitochondria (Fig. 7). Newly synthesized Pex15 is targeted to peroxisomes, presumably directly, in a manner dependent on Pex19 and Pex3, but independently of the GET pathway. When Pex15 is mistargeted to the mitochondrial OM due to the loss of Pex19 or Pex3, it is recognized by Msp1 and extracted from the mitochondrial OM into the cytosol. After extraction, Pex15 is either transferred to the ER *via* the GET pathway or to peroxisomes *via* a Pex19- and Pex3-dependent pathway. The Pex15 transferred to the ER is recognized by the E3 ligases Doa10 and Hrd1, which target it for proteasomal degradation. However, the Pex15 on the peroxisomal membrane is also recognized by the Msp1 on peroxisomes and extracted into the cytosol, but can be returned to peroxisomes with the assistance of Pex19 and Pex3.

**Figure 7.**
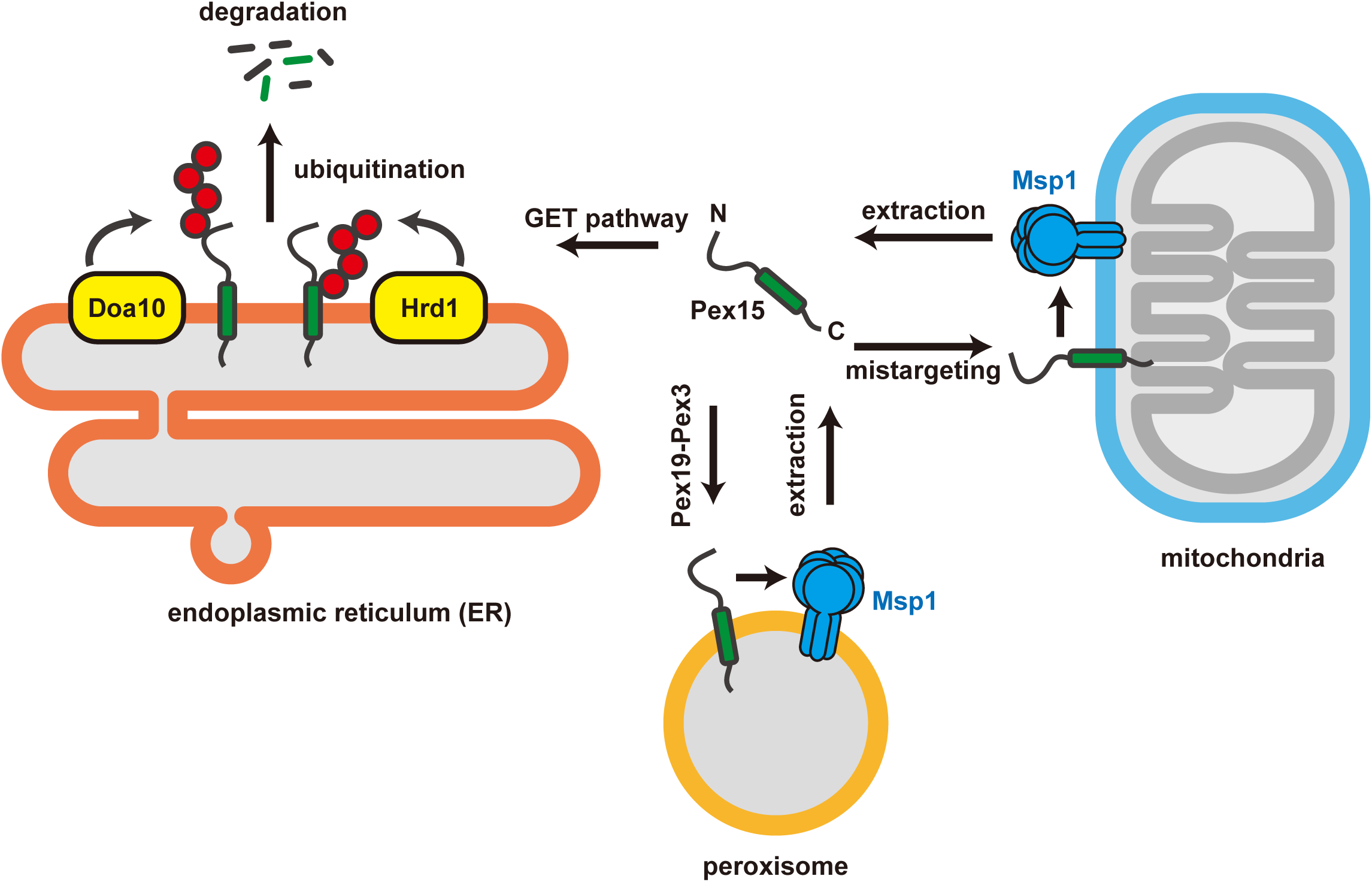
Model of the Msp1- and Pex19- and Pex3-mediated localization of Pex15 to peroxisomes. After translation, newly synthesized Pex15 binds to Pex19 in the cytosol and is directed to the peroxisomal membrane. Membrane insertion occurs through the association of Pex19 with Pex3. The absence of Pex19 or Pex3 results in the mistargeting of Pex15 to the mitochondrial OM. Mislocalized Pex15 on the mitochondrial OM is extracted by Msp1 and transferred to the ER *via* the GET pathway. At the ER, Pex15 is ubiquitinated by the ER-resident E3 ligase Doa10 or Hrd1 and subsequently degraded by the proteasome. In contrast, when Pex19 and Pex3 are present, Msp1-extracted Pex15 from the mitochondrial OM can be correctly re-directed to peroxisomes. In addition, Pex15 is occasionally extracted from the peroxisomal membrane by Msp1, but is reinserted into peroxisomes in a Pex19- and Pex3-dependent manner.

Yeast Pex15 consists of 383 amino acids, with an N-terminal cytosolic domain, a TM segment (residues 335-349), and a C-terminal tail (residues 353-383). Previously, Pex19 was shown to bind to both the TM segment and C-terminal tail of Pex15, and the fusion of the Pex19 binding site was sufficient for peroxisomal targeting (Halbach et al., 2006). The GET pathway was proposed to play a role in targeting Pex15 to the ER, followed by its transfer to peroxisomes *via* vesicular transport (Schuldiner et al., 2008). Several studies have supported an ER-mediated, indirect transport pathway for Pex15 (van der Zand et al., 2010; Lam et al., 2010; Chen et al., 2014; Okreglak and Walter, 2014); for example, overexpressed Pex15 from the *GAL1* promoter is mislocalized to mitochondria in yeast mutants with a defective GET pathway. In contrast to these results, another study reported that Pex15, when expressed from its endogenous promoter, can still localize to peroxisomes even in GET pathway-deficient cells (Li et al., 2019). Consistent with this finding, our results demonstrated that Pex15 expressed from its native chromosomal locus localizes to peroxisomes, despite mutations in the GET pathway (Fig. 1A). Although earlier studies suggested that Pex19 and Pex3 facilitate the transfer of Pex15 from the ER to peroxisomes *via* vesicular transport, our results show that the peroxisomal targeting of newly synthesized Pex15 is significantly suppressed by the acute depletion of AID-tagged Pex19 (Figs. 3D, E) and Pex3 (Figs. 3G, H). Additionally, the depletion of Pex19 causes Pex15 mistargeting to mitochondria, rather than the ER, especially in *MSP1*-deficient cells (Fig. S3A-D). Furthermore, the GET pathway is not required for the peroxisomal targeting of Pex15 (Figs. S3E, F). In conclusion, the peroxisomal targeting of Pex15 occurs primarily through a Pex19- and Pex3-dependent pathway, with no involvement of the GET pathway. Instead, the GET pathway appears necessary for moving the mistargeted Pex15 from mitochondria to the ER. However, we cannot completely exclude the possibility that Pex19 and Pex3 are partly involved in sorting Pex15 from the ER to peroxisomes *via* vesicular transport.

We have provided evidence that the Msp1 on the mitochondrial OM cooperates with Pex19 in the transfer of Pex15 from mitochondria to peroxisomes. Specifically, even when Pex15 is mistargeted to the mitochondria, it can still bind to Pex19 after being extracted into the cytosol by Msp1. Once Pex15 is loaded onto Pex19, it is re-directed to Pex3 on the peroxisomal membrane (Fig. 4). This suggests that Msp1 can proofread the mislocalization of other TA proteins, beyond those delivered *via* the GET pathway. However, once Pex15 is mistargeted to the mitochondrial OM, there could be competition between its peroxisomal targeting *via* the Pex19-Pex3-dependent pathway and its ER targeting *via* the GET pathway (Fig. S4). This is because Pex19 and Get3 compete for binding to the Pex15 in the cytosol after its extraction by Msp1. Under our experimental conditions, Pex15 is overexpressed and exceeds the binding capacity of Pex19, which may lead to the excess Pex15 being targeted for degradation in the ER (Fig. S4).

Endogenous Pex15 on peroxisomes remains stable in WT cells (Fig. 1A). A model to explain this stability suggests that the Pex15 on peroxisomes binds to Pex3, thereby preventing its recognition by peroxisomal Msp1 (Weir et al., 2017). Weir et al. demonstrated that the fluorescent signal of Pex15 decreased in an Msp1-dependent manner when Pex3, fused with an auxin degron tag (Pex3-AID), was rapidly depleted by auxin treatment. Our results show that the rapid degradation of AID-tagged Pex19 leads to the Msp1-dependent degradation of the endogenous Pex15 on peroxisomes (Figs. 5A, B). Our data indicate that, upon rapid depletion of Pex3-AID, Pex15 is extracted from peroxisomes by Msp1 but remains stable, presumably in the cytosol (Fig. S5A, B). Furthermore, the endogenous Pex3 remains stable even after the rapid degradation of AID-Pex19 (Fig. S5C). We suggest that the previously proposed model alone does not fully explain the stability of Pex15 on peroxisomes. Given the involvement of Pex19 and Pex3 in the targeting of Pex15 to peroxisomes, we propose an alternative dynamic localization mechanism. In this model, the Msp1 on peroxisomes constantly extracts Pex15 from the membrane, while Pex19 protects the extracted Pex15 from degradation in the cytosol and facilitates its return to the peroxisomes (Fig. 7).

## Materials and methods

### Strains and plasmids

The yeast strains and plasmids used in this study are described in Tables S1 and S2, respectively. Yeast strains were constructed using standard homologous recombination methods (Knop et al., 1999).

For the fluorescence microscopy imaging in Figs. 1, 2, 5, 6, S1 and S5, the mNeonGreen (mNG) tag was integrated immediately after the start codon of the *PEX15* ORF by using CRISPR-Cas9 genome editing derived from *Streptococcus pyogenes*, according to the published protocol (Okada et al., 2021). For the fluorescence microscopy imaging in Fig. S1, the mNG tag was integrated into the position following the start codon of the *PEX19* ORF by using CRISPR-Cas9 genome editing derived from *Staphylococcus aureus*, according to the published protocol (Okada et al., 2021). For ER (pMT328), mitochondria (pMT367) and peroxisome (pMT375) labeling, we linearized the expression plasmids by an inverse PCR reaction and integrated them into the *URA3* locus of the yeast chromosome.

For the cycloheximide (CHX) chase experiments in Figs. 1, 2, 5, S2 and S5, the 3xFLAG tag was integrated into the position following the start codon of the *PEX15* ORF, using the same method as for the mNG-tag insertion.

For the fluorescence microscopy imaging in Figs. 3 and S3, OsTIR1-9xMyc was expressed from the *ADH1* promoter by integrating the linearized pSMQ249 into the *LEU2* locus (Nishimura et al., 2009). Subsequently, the AID*(IAA17^71-114^)-tag was integrated into the position following the start codon of the *PEX19* ORF in the OsTIR1-expressing yeast cells, by using CRISPR-Cas9 genome editing derived from *S. aureus*. The AID* -V5 tag was integrated into the position preceding the stop codon of the *PEX3* ORF in the OsTIR1-expressing yeast cells. 3xFLAG-mNG-Pex15 was expressed from the *DDI2* promoter by integrating the linearized pSMQ175 into the *TRP1* locus in AID*-Pex19- and Pex3-AID*-V5-expressing yeast cells. Pex11-mCherry or Tom70-mCherry was expressed from its own promoter by integrating the mCherry tag (pMT294) into the position preceding the stop codon of the *PEX11* or *TOM70* ORF, respectively.

For the fluorescence microscopy imaging in Fig. 4, the expression cassette for Z4EV (pSMQ445) was integrated into the *HIS3* locus in AID*-Pex19-expressing yeast cells. We deleted the *MSP1* gene, since a fraction of Msp1 is targeted to peroxisomes. We then expressed the Msp1 variant exclusively targeted to mitochondria by replacing the N-terminal TM segment of Msp1 with that of Tom70 (Tom70TM-Msp1) in the AID*-Pex19-expressing yeast cells. Tom70TM-Msp1 was expressed from the β-Estradiol *Z4EV* promoter by integrating the linearized pSMQ451 into the *URA3* locus in the AID*-Pex19-expressing yeast cells.

For the fluorescence microscopy imaging in Fig. 6, Tom70TM-Msp1-V5 or Pex22TM-Msp1-V5 was expressed from its own promoter by integrating the linearized pSMQ490 or pSMQ485 into the *URA3* locus in *msp1*11 cells expressing mNG-Pex15. All plasmid constructs were verified by DNA sequencing.

### Yeast growth conditions

Yeast strains were cultured in YP (1% yeast extract and 2% polypeptone) or SC (0.67% yeast nitrogen base without amino acids and 0.5% casamino acids) media with appropriate carbon sources (2% glucose (D)) and supplements.

### Cycloheximide chase

Yeast cells expressing 3xFLAG-Pex15 under the control of its own promoter from the chromosome were grown in YPD medium at 30°C. After an overnight incubation at 30°C, the cells were diluted to an OD_600_ of 0.2-0.3 and grown in YPD medium at 30°C. When the OD_600_ reached 1.0, cycloheximide (CHX) was added at a final concentration of 100 μg/ml to start a chase at 30°C. One OD_600_ unit of cells was collected periodically and frozen at -80°C until use. The frozen cells were thawed in ice-cold 0.1 M NaOH and incubated on ice for 5 min. The cell pellets were precipitated by centrifugation at 20,000 × *g* for 5 min. The resulting pellet was resuspended in SDS-PAGE sample buffer, and analyzed by SDS-PAGE and immunoblotting.

### Immunoblotting

Proteins were transferred from polyacrylamide gels to PVDF membranes (Bio-Rad) in blotting buffer for SDS-PAGE (25 mM Tris, 192 mM glycine, 5% methanol), at a constant voltage of 25 V. After transfer, the membranes were washed in TBST buffer (10 mM Tris-HCl, pH 7.5, 150 mM NaCl, 0.05% Tween-20) and blocked with 5% skim milk in TBST buffer for 1 h. The membranes were incubated with primary antibodies in TBST buffer overnight at 4°C and washed 3 times with TBST (5 min for each wash). The membranes were then incubated with secondary antibodies for 30 min and washed 3 times with TBST. The PVDF membranes were scanned using an Amersham Typhoon scanner (Cytiva), and signal intensities were quantified by using the ImageQuant software (Cytiva).

We used the following antibodies: monoclonal anti-FLAG M2 mouse IgG (F3165; 1:2,000; Millipore-Sigma), monoclonal anti-Pgk1 mouse IgG (ab113687; 1:2,000; Abcam), monoclonal anti-V5 mouse IgG (M167-3; 1:2,000; Medical & Biological Laboratories), polyclonal anti-Tom40 rabbit IgG (1:2,000), polyclonal anti-Msp1 rabbit IgG (1:2,000), polyclonal anti-Pex19 rabbit IgG (1:2,000), Goat anti-Mouse IgG (H+L) Secondary Antibody, DyLight 800 4X PEG (SA5-35521; 1:2,000; Thermo Fisher Scientific), Goat anti-Rabbit IgG (H+L) Secondary Antibody, DyLight 800 4X PEG (SA5-35571; 1:2,000; Thermo Fisher Scientific), Goat anti-Mouse IgG (H+L) Cross-Adsorbed Secondary Antibody, Cyanine5 (A10524; 1:2,000; Thermo Fisher Scientific), and Goat anti-Rabbit IgG (H+L) Cross-Adsorbed Secondary Antibody, Cyanine5 (A10524; 1:2,000; Thermo Fisher Scientific).

### Fluorescence microscopy

Yeast cells were mounted on a 1% agarose pad containing TBS buffer, and were placed on a 14-mm glass bottom dish (Matsunami Glass). Live cell imaging in this study was performed using a THUNDER Ready DMi8 microscope (Leica Microsystems), equipped with an HCX PL APO 100×/1.40–0.70 OIL objective lens (Leica Microsystems), a K8 Scientific CMOS camera (Leica Microsystems), and an LED3 light source (Leica Microsystems). Fluorescence images were processed with the THUNDER imaging system (Leica Microsystems) with the following settings. Live cell images obtained with the THUNDER Ready DMi8 microscope at room temperature were processed using the THUNDER Imaging System (Leica Microsystems), using the Large Volume Computational Clearing (LVCC) mode. The z-sections were collected every 0.2 - 0.3 μm from the top to bottom surface of the yeast cells. Acquired images were processed and analyzed with the Fiji software.

For the live cell imaging in Figs. 1A, 1C, 2B, 2C, 2F, 6B and S1, yeast cells were grown in SCD medium. After an overnight incubation at 30°C, the cells were diluted to an OD_600_ of 0.2-0.3 and grown in SCD medium at 30°C. When the OD_600_ reached 0.8-1.0, live cell imaging was performed using the THUNDER Ready DMi8 microscope.

For the Turn-ON/OFF system using the AID system in Figs. 3D, 3G, S3A, S3C and S3E, yeast cells were grown in SCD medium. After an overnight incubation at 30°C, the cells were diluted to an OD_600_ of 0.2-0.3 and grown in SCD medium at 30°C. When the OD_600_ reached 0.8-1.0, the cells were treated with or without 1 mM IAA for 30 min at 30°C in SCD medium, and then further incubated for 2 h at 30°C in the presence of 5 mM cyanamide to induce 3xFLAG-mNG-Pex15 expression from the *DDI2* promoter. Live cell imaging was performed using the THUNDER Ready DMi8 microscope.

For the Msp1-mediated transfer of mislocalized Pex15 using the AID system in Figs. 4C-E, yeast cells were grown in SCD medium. After an overnight incubation at 30°C, the cells were diluted to an OD_600_ of 0.2-0.3 and grown in SCD medium at 30°C. When the OD_600_ reached 0.8-1.0, the cells were treated with 1 mM IAA for 30 min at 30°C in SCD medium, and then further incubated for 2 h at 30°C in the presence of 5 mM cyanamide to induce 3xFLAG-mNG-Pex15 expression from the *DDI2* promoter. After the expression of 3xFLAG-mNG-Pex15, the cells were washed with fresh SCD medium twice and incubated for 2 h at 30°C. Time-lapse imaging using the THUNDER Ready DMi8 microscope was performed after the addition of 1 μM β-Estradiol to start the expression of Tom70TM-Msp1.

For the cycloheximide chase experiment in Figs. 5B-C and S5B, yeast cells were grown in SCD medium at 30°C. After an overnight incubation at 30°C, the cells were diluted to an OD_600_ of 0.2-0.3 and grown in SCD medium at 30°C. When the OD_600_ reached 0.8-1.0, the cells were treated with 1 mM IAA and 100 μg/ml CHX, followed by live cell imaging using the THUNDER Ready DMi8 microscope every 30 minutes.

### Statistics

Statistical tests used for data analysis are defined in the figure legends. These include one-way ANOVA with Dunnett’s multiple comparison test in Fig. 2F, Fig. 4F, and Fig. 6C, and the two-tailed paired *t* test in Fig. 3E, H; and Fig. S3B, D, F. Data distribution was assumed to be normal, but this was not formally tested.

## Online supplemental material

Fig. S1 shows the localization of mNG-Pex19 in WT and *pex3*11 cells. Fig. S2 shows the CHX chase experiment of 3xFLAG-Pex15 in *pex19*11*pdr5*11 cells, with and without MG132 treatment. Fig. S3 shows the analysis of peroxisomal targeting of 3xFLAG-mNG-Pex15 in AID*-Pex19 *msp1*11 cells and Get3-AID*-9xMyc cells. Fig. S4 shows that the Pex15 extracted from the mitochondrial OM by Msp1 can be targeted to the ER *via* the GET pathway for degradation. Fig. S5 shows that endogenous Pex15 is extracted by peroxisomal Msp1 under Pex3-AID*-V5 depleted conditions. Online material is available at http://ww.jcb.org/cgi/content/full/…..

## Acknowledgements

We thank the members of the Numata and Endo labs for discussions and critical comments on the manuscript. We are grateful to the Center for Advanced Instrumental and Educated Support at the Faculty of Agriculture, Kyushu University, for technical assistance with fluorescence microscopy.

This work was supported by JSPS KAKENHI grants to S.M. (20H05929, 23H02437 and 23K17995), T.E. (15H05705, 20H04912, 20H00458, 20H05689, and 20H05929), and S.O. (24KJ0210 and 24K18117), an AMED-CREST grant to T.E. (21gm1410002), and a grant from the Japan Foundation for Applied Enzymology to S.M., and a grant from Kinoshita Foundation to S.M., and a grant from Takeda Science Foundation to S.M. and T.E..

The authors declare no competing financial interests.

## Author contributions

Conceptualization, methodology, project administration, supervision, and writing the original draft were accomplished by S. Matsumoto and T. Endo. T. Numata contributed to the project administration and supervision. The investigation was primarily carried out by S. Matsumoto, and partly by Y. Kogure and S. Ono. S. Matsumoto and Y. Kogure (in part) were responsible for data curation, formal analysis, and visualization.

---

**Figure S1.**
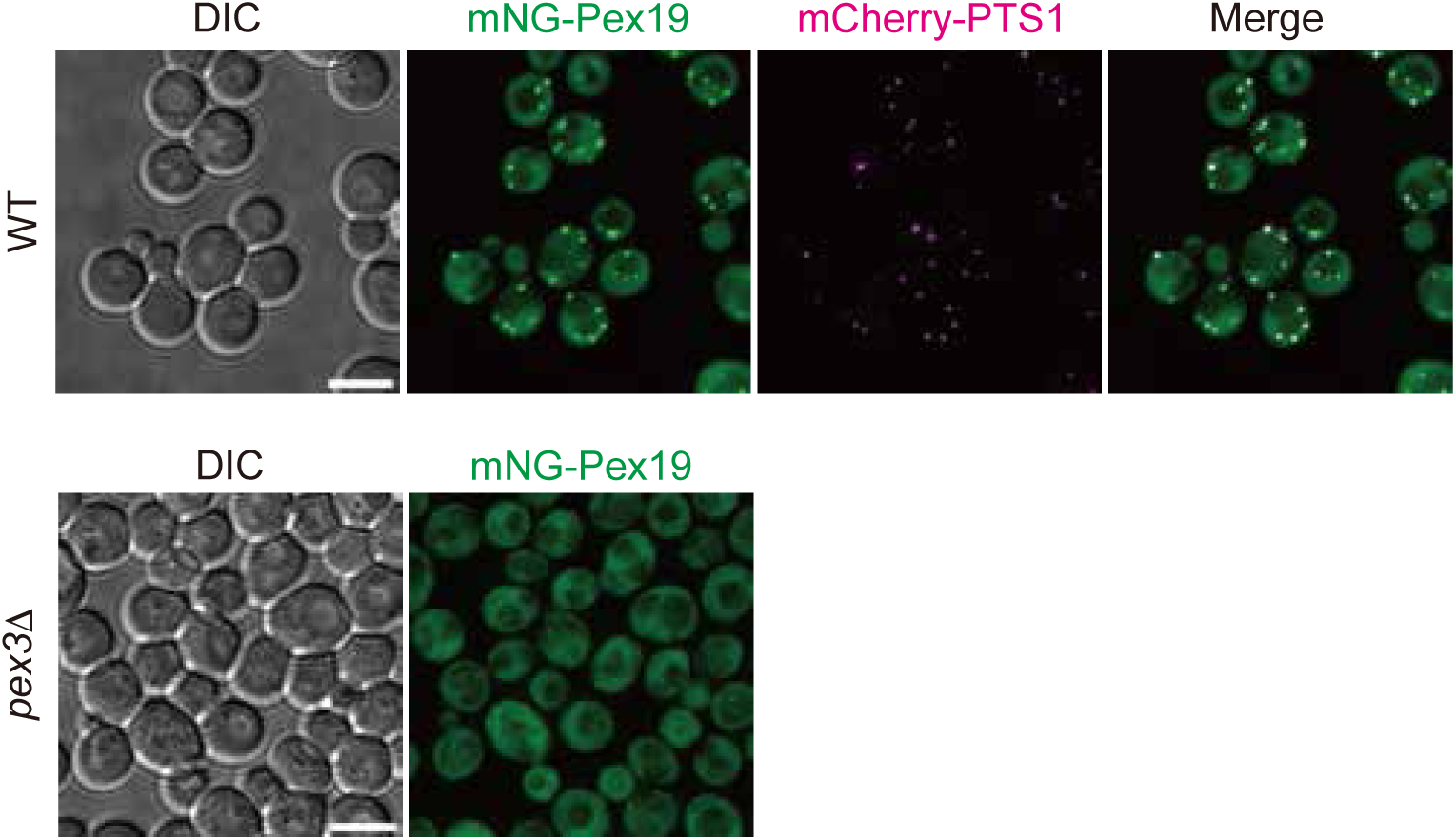
Localization of mNG-tagged Pex19 in WT and pex3Δ cells. WT or *pex3*11 cells expressing mNG-Pex19 under the control of its own promoter from the chromosome were grown in SCD medium at 30°C and imaged by fluorescence microscopy. Peroxisomes were labeled with mCherry-PTS1. Maximum projection images of Z-stacks are shown. Scale bar, 5 μm. DIC, differential interference contrast microscopy.

**Figure S2.**
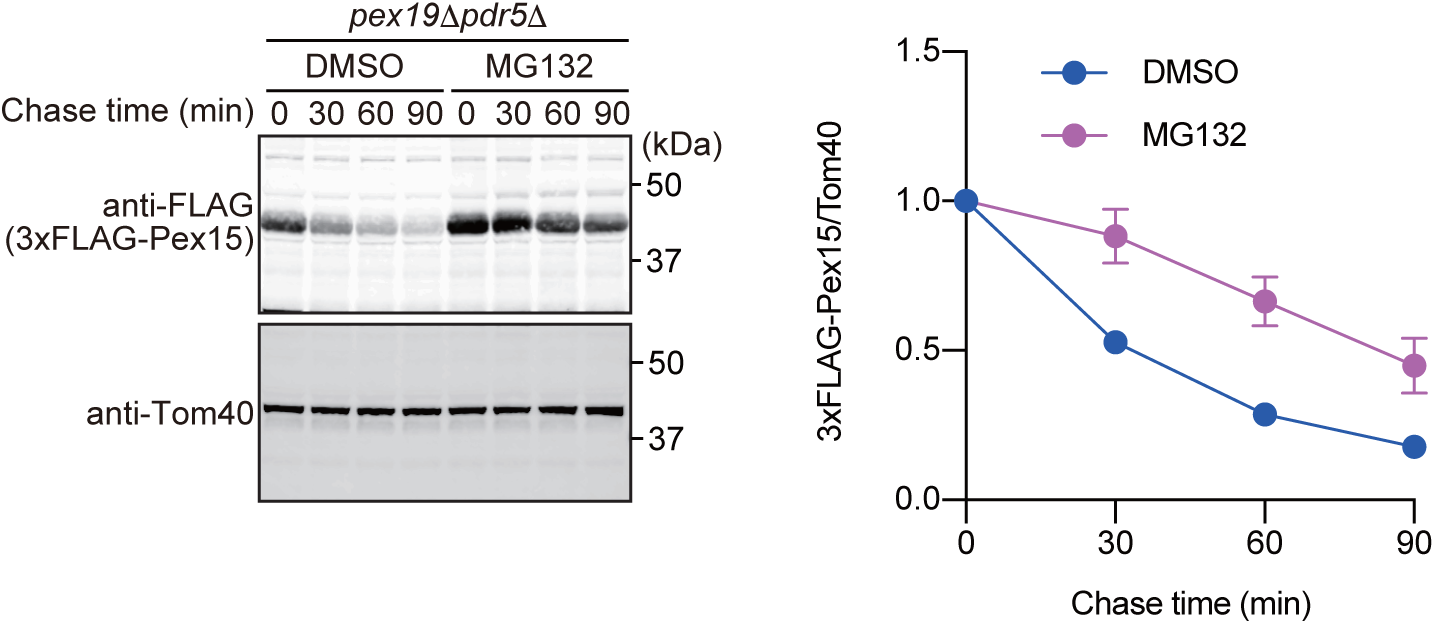
Proteasome is involved in the degradation of mislocalized Pex15. *pex19*11*pdr5*11 cells expressing 3xFLAG-Pex15 under the control of its own promoters from the chromosomes were grown in YPD medium at 30°C, and then further incubated for 30 min after the addition of 50 μM MG132. Extracts were prepared at the indicated times from cells incubated at 30°C after the addition of 100 µg/ml CHX, and proteins were analyzed by SDS-PAGE and immunoblotting with the indicated antibodies (left). The normalized relative amounts of 3xFLAG-Pex15 were plotted against the chase time. Values are the mean ± S.D. from three independent experiments.

**Figure S3.**
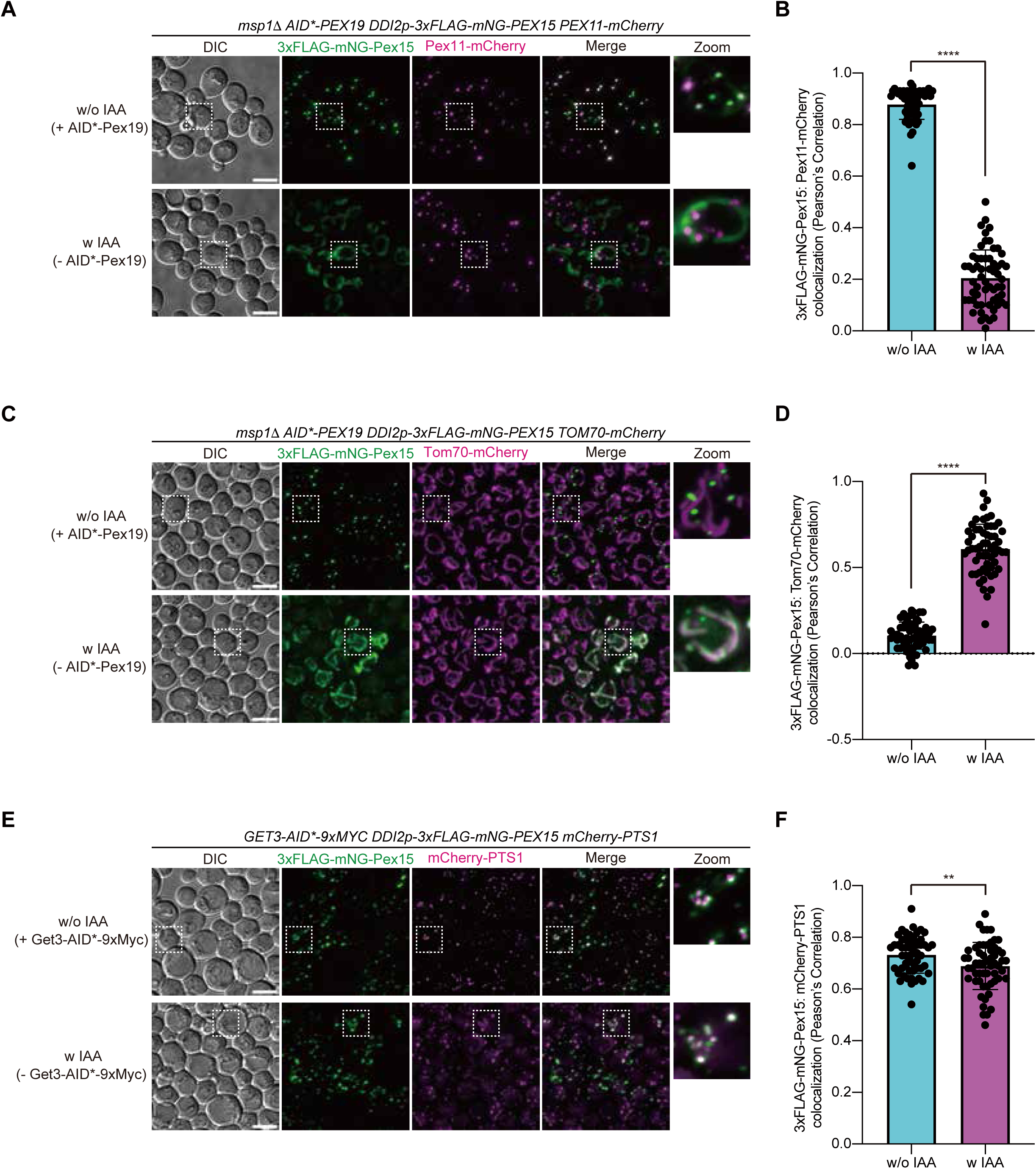
Newly synthesized Pex15 is mistargeted to mitochondria by rapid depletion of Pex19 in *MSP1*-deficient cells. (**A**) *msp1*11 cells expressing AID*-Pex19 under the control of its own promoter from the chromosome were grown in SCD medium at 30°C. AID*-Pex19 was depleted by a treatment with 1 mM IAA for 30 min at 30°C, followed by 3xFLAG-mNG-Pex15 induction with 5 mM cyanamide at 30°C for 2 hours. The localization of 3xFLAG-mNG-Pex15 was imaged by fluorescence microscopy in the presence (-IAA, upper panel) or absence (+IAA, lower panel) of AID*-Pex19. Maximum projection images of Z-stacks are shown. Peroxisomes were labeled with Pex11-mCherry. Scale bar, 5 μm. DIC, differential interference contrast microscopy. (**B**) Colocalization of 3xFLAG-mNG-Pex15 with peroxisomes was analyzed using Pearson’s correlation coefficient between the mNG and mCherry signals. Values are the mean ± S.D. (-IAA; *n*=60, +IAA; *n*=60) from three independent replicates; *n* represents the number of cells. ****, P < 0.0001 compared with IAA untreated (-IAA) cells by two-tailed paired *t* test. **(C)** The localization of 3xFLAG-mNG-Pex15 was imaged by fluorescence microscopy in the presence (-IAA, upper panel) or absence (+IAA, lower panel) of AID*-Pex19, as in (A). Maximum projection images are shown. Mitochondria were labeled with Tom70-mCherry. **(D)** Colocalization of 3xFLAG-mNG-Pex15 with mitochondria was analyzed using Pearson’s correlation coefficient between the mNG and mCherry signals, as in (B). Values are the mean ± S.D. (-IAA; *n*=59, +IAA; *n*=58) from three independent replicates; *n* represents the number of cells. ****, P < 0.0001 compared with IAA untreated (-IAA) cells by two-tailed paired *t* test. (**E**) Yeast cells expressing Get3-AID*-9xMyc under the control of its own promoter from the chromosome were grown in SCD medium at 30°C. Get3-AID*-9xMyc was depleted by treatment with 1 mM IAA for 30 min at 30°C, followed by induction of 3xFLAG-mNG-Pex15 with 5 mM cyanamide at 30°C for 2 hours. Localization of 3xFLAG-mNG-Pex15 was imaged by fluorescence microscopy in the presence (-IAA, upper panel) or absence (+IAA, lower panel) of Get3-AID*-9xMyc. Maximum projection images are shown. Peroxisomes were labeled with mCherry-PTS1. (**F**) Colocalization of 3xFLAG-mNG-Pex15 with peroxisomes was analyzed using Pearson’s correlation coefficient between the mNG and mCherry signals, as in (B). Values are the mean ± S.D. (-IAA; *n*=56, +IAA; *n*=57) from three independent replicates; *n* represents the number of cells. **, P = 0.0061 compared with IAA untreated (-IAA) cells by two-tailed paired *t* test.

**Figure S4.**
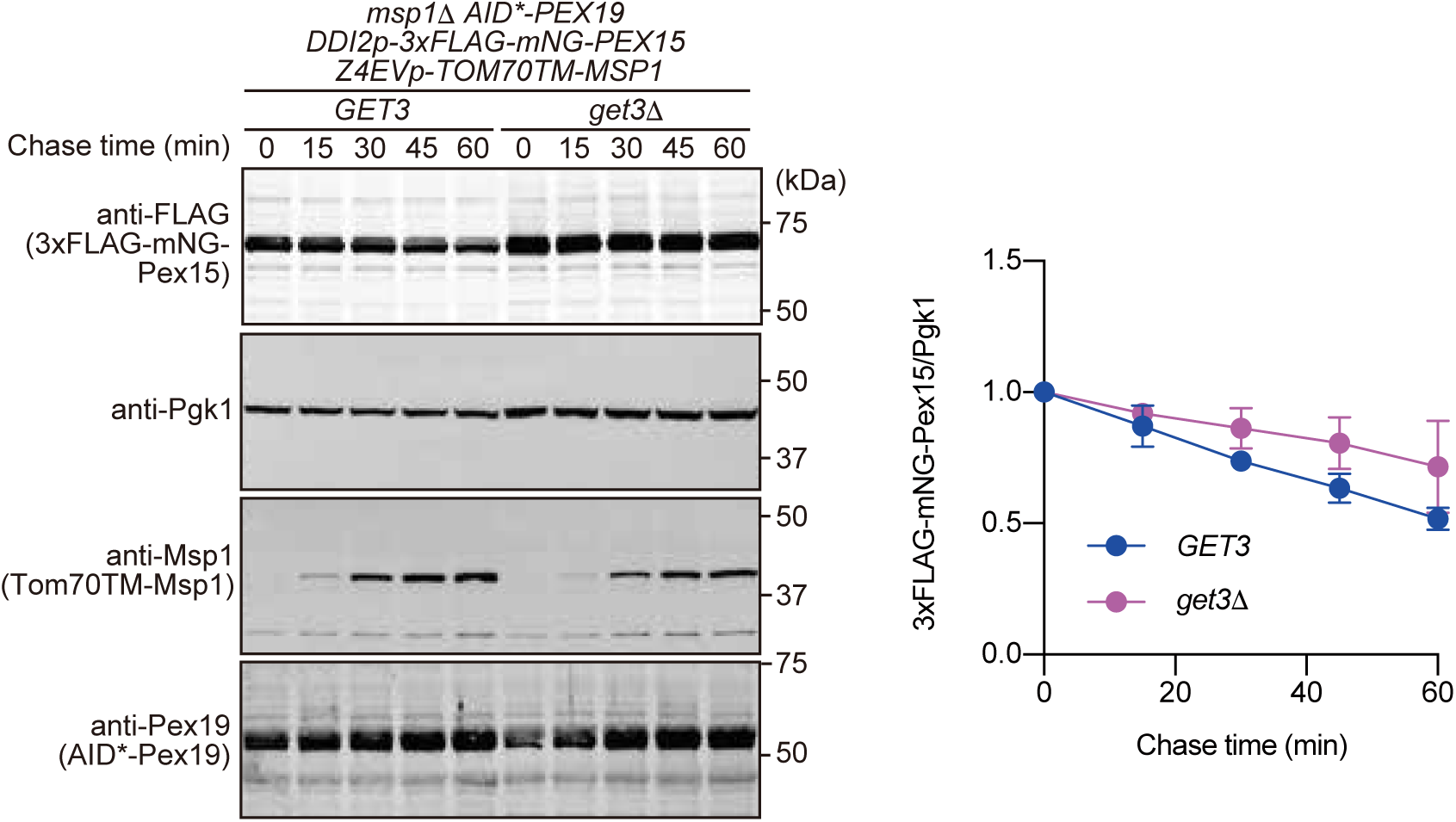
The GET pathway mediates Pex15 transfer from mitochondria to the ER for degradation. The promoter shutoff chase of 3xFLAG-mNG-Pex15 expression was performed with the yeast cells described in Fig. 4A (*GET3*) and yeast cells lacking *GET3* (*get3*11), which were grown in SCD medium at 30°C. AID*-Pex19 was then depleted by a treatment with 1 mM IAA for 30 min at 30°C, followed by the induction of 3xFLAG-mNG-Pex15 with 5 mM cyanamide for 2 h at 30°C. The cells were washed with fresh SCD medium to shut off the 3xFLAG-mNG-Pex15 expression and induce the re-expression of AID*-Pex19 by an incubation in fresh SCD medium for 2 h at 30°C. Cell extracts were prepared at the indicated times in the presence (+β-Estradiol) of Tom70TM-Msp1 induction, and proteins were analyzed by SDS-PAGE and immunoblotting with the indicated antibodies. The normalized relative amounts of 3xFLAG-mNG-Pex15 to Pgk1 were plotted against the chase time. Values are the mean ± S.D. from three independent experiments.

**Figure S5.**
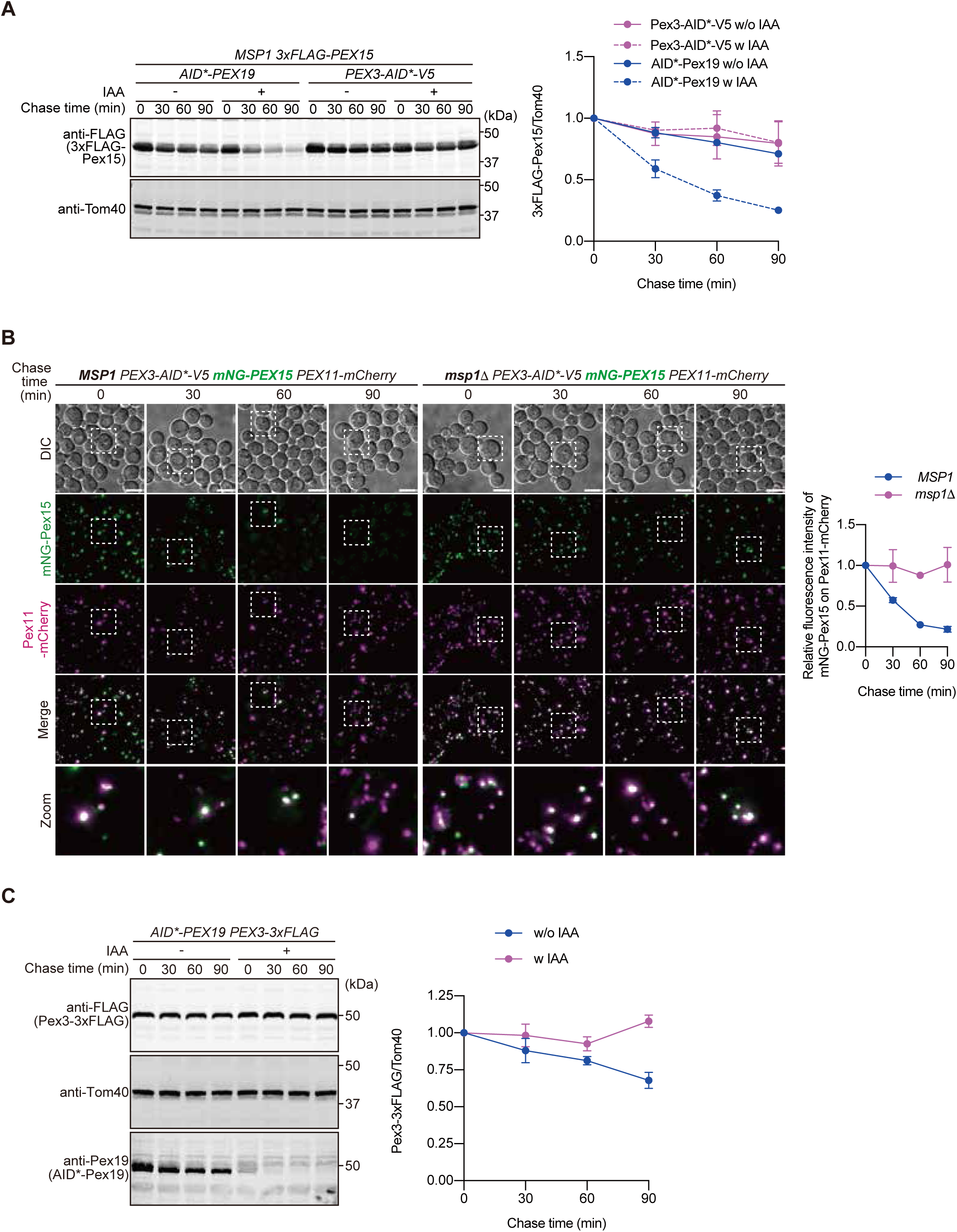
Peroxisomal Msp1 extracts endogenous Pex15 upon rapid depletion of Pex3. **(A)** Yeast cells expressing 3xFLAG-Pex15 with AID*-Pex19 or Pex3-AID*-V5 under the control of their own promoters from the chromosomes were grown in YPD medium at 30°C, and then further incubated for 30 min after the addition of 1 mM IAA. Extracts were prepared at the indicated times from cells incubated at 30°C after the addition of 100 µg/ml CHX, and proteins were analyzed by SDS-PAGE and immunoblotting with the indicated antibodies (left). The normalized relative amounts of 3xFLAG-Pex15 were plotted against the chase time. Values are the mean ± S.D. from three independent experiments. **(B)** Wild-type (*MSP1*) and *msp1*11 cells expressing Pex3-AID*-V5 and mNG-Pex15 under the control of their own promoters from the chromosomes were grown in SCD medium at 30°C. Localization of mNG-Pex15 was observed by fluorescence microscopy at the indicated time points after the addition of 1 mM IAA and 100 µg/ml CHX. Maximum projection images of Z-stacks are shown. Peroxisomes were labeled with Pex11-mCherry. Scale bar, 5 μm. DIC, differential interference contrast microscopy. The quantification of the mNG-Pex15 signal was performed as in Fig. 5B, with the signal at time 0 set to 1, and the average values obtained from two independent experiments were plotted. **(C)** Yeast cells expressing AID*-Pex19 and Pex3-3xFLAG under the control of their own promoters from the chromosomes were grown in YPD medium at 30°C, and were further incubated for 30 min after the addition of 1 mM IAA. Extracts were prepared at the indicated times from cells incubated at 30°C after the addition of 100 µg/ml CHX, and proteins were analyzed by SDS-PAGE and immunoblotting with the indicated antibodies (left). The normalized relative amounts of Pex3-3xFLAG were plotted against the chase time. Values are the mean ± S.D. from three independent experiments.

